# A sporadic Alzheimer’s blood-brain barrier model for developing ultrasound-mediated delivery of Aducanumab and anti-Tau antibodies

**DOI:** 10.1101/2022.03.06.483200

**Authors:** Joanna M. Wasielewska, Juliana C. S. Chaves, Rebecca L. Johnston, Laura A. Milton, Damián Hernández, Liyu Chen, Jae Song, Wendy Lee, Gerhard Leinenga, Rebecca M. Nisbet, Alice Pébay, Jürgen Götz, Anthony R. White, Lotta E. Oikari

## Abstract

**Rationale:** The blood-brain barrier (BBB) is a major impediment to therapeutic intracranial drug delivery for the treatment of neurodegenerative diseases, including Alzheimer’s disease (AD). Focused ultrasound applied together with microbubbles (FUS^+MB^) is a novel technique to transiently open the BBB and increase drug delivery. Evidence suggests that FUS^+MB^ is safe, however the effects of FUS^+MB^ on human BBB cells, especially in the context of AD, remain sparsely investigated.

**Methods:** Here we generated BBB cells (induced brain endothelial cells (iBECs) and astrocytes (iAstrocytes)) from apolipoprotein E gene allele E4 (*APOE4*, high AD risk) and allele E3 (*APOE3*, lower AD risk) carrying patient-derived induced pluripotent stem cells (iPSCs). We then developed a human sporadic AD BBB cell platform to investigate the effects of FUS^+MB^ on BBB cells and screen for the delivery of two potentially therapeutic AD antibodies.

**Results:** We utilized this robust and reproducible human BBB model to demonstrate increased delivery of therapeutic AD antibodies across the BBB following FUS^+MB^ treatment, including an analogue of Aducanumab (Aduhelm^TM^; anti-amyloid-β) and a novel anti-Tau antibody RNF5. Our results also demonstrate the safety of FUS^+MB^ indicated by minimal changes in the cell transcriptome as well as little or no changes in cell viability and inflammatory responses within the first 24 h post FUS^+MB^. Finally, we report a more physiologically relevant hydrogel-based 2.5D BBB model as a key development for FUS^+MB^-mediated drug delivery screening, with potentially higher translational utility.

**Conclusion:** Our results demonstrate an important translatable patient BBB cell model for identifying FUS^+MB^-deliverable drugs and screening for cell- and patient-specific effects of FUS^+MB^, accelerating the use of FUS^+MB^ as a therapeutic modality in AD.

**One Sentence Summary:** Focused ultrasound increases the *in vitro* delivery of therapeutic antibodies Aducanumab and anti-Tau in a sporadic Alzheimer’s disease patient-derived blood-brain barrier cell model.

## INTRODUCTION

Alzheimer’s disease (AD) is the most prevalent cause of dementia characterized by progressive and irreversible cognitive decline, including loss of short- and long-term memory [1]. The majority of AD cases (>95%) are described as sporadic with late disease on-set (over 65 years of age) and a genetic etiology that is not fully understood. However, several genetic risk factors have been identified for sporadic AD, including the E4 allele of the apolipoprotein E (*APOE)* gene [2, 3]. Relative to the most common *APOE E3/E3* isoform, the presence of *APOE E4* allele confers an increased risk of developing AD [2, 3]. The most commonly described pathological hallmarks of AD include the accumulation of amyloid-β plaques (Aβ) and phosphorylated-Tau (p-Tau) containing neurofibrillary tangles [1, 4, 5]. However, emerging evidence indicates that changes in the brain vasculature are also associated with AD [6].

The blood-brain-barrier (BBB) is a selectively permeable barrier at the blood-brain interface, formed by brain endothelial cells (BECs) and supporting cells, including astrocytes and pericytes, and its role is to maintain a highly controlled internal brain milieu [7]. High integrity of the BEC monolayer is essential for barrier function of the BBB and is achieved by the high expression of tight junction (TJ) and adherens junction (AJ) proteins between adjacent BECs, including occludin, claudin-5, ZO-1 and vascular endothelial (VE)-cadherin respectively [8]. Synergistically, TJ and AJ proteins ensure a high trans-endothelial electrical resistance (TEER) of the BBB and limit paracellular permeability through the BBB into the brain [7, 9, 10]. Recently, the association of the *APOE4* isoform with BBB breakdown has been reported [11]. It has been suggested that BBB dysfunction in *APOE4* carriers occurs before cognitive decline and is independent of Aβ and p-Tau accumulation, and thus an additional driving force of cognitive impairment [11]. Studies in transgenic mice further supported a role for *APOE4* in BBB dysfunction, reflected by an early BBB breakdown and cerebral microhemorrhages, which lead to secondary neurodegeneration [8, 11-13].

The BBB is not only altered in AD, but it also poses a major physical obstacle for drug delivery into the brain. The BBB actively restricts the entry of over 98% of small molecule drugs and up to almost 100% of large therapeutic antibodies and growth factors into the brain, presenting a key hurdle in successful AD drug discovery [14]. Furthermore, the altered brain milieu caused by BBB disruption in AD likely further hinders controlled drug delivery and metabolization [6], posing an urgent need for adequate AD *in vitro* and preclinical models to understand AD-specific changes in the BBB.

With a concerning 99.6% failure rate of clinical trials targeting AD, a paradigm-shift may originate from the development of accurate human-like pre-clinical models of drug delivery that improve the penetration of therapeutics through the BBB [15]. Correspondingly, a recent study by Ohshima *et al.* compared the permeability of several drugs of known pharmacokinetics in various *in vitro* BBB models, showing that human induced pluripotent stem cell (hiPSC)-derived BBB models correlated most closely with human *in vivo* BBB permeability [16]. This suggests that patient-derived hiPSC-based BBB models provide a clinically relevant platform for permeability studies in AD drug discovery.

Focused low-intensity ultrasound (FUS) applied together with microbubbles (MBs) is emerging as a novel technique to transiently open the BBB to aid in drug delivery into the brain. The ultrasound wave causes intravenously injected MBs to expand and contract, and when in contact with the BBB, exert a force that results in the loosening of tight junctions leading to transient BBB opening [17]. We recently demonstrated the ability to model FUS-mediated drug delivery and Aβ clearance *in vitro* using a familial AD iPSC-derived induced brain endothelial cell-like (iBEC) model [18]. Our findings reported differences in immediate and long-term responses to FUS^+MB^ in familial AD compared to control iBECs, suggesting disease- or patient-specific effects. Importantly, the safety of ultrasound-mediated BBB opening has been demonstrated in multiple animal and several small patient studies, although drug delivery in AD patients has not yet been achieved [19-22]. As a detailed understanding of the mechanisms and long-term safety of FUS-mediated human-BBB opening and its secondary effects is still largely needed [23], a patient cell platform to study the effects of FUS and identify FUS-deliverable drugs, especially for sporadic AD, would be an important step in improving and tailoring FUS-therapies for AD patients.

Here we developed a sporadic AD patient-derived BBB-like *in vitro* model to study the effects of FUS^+MB^ on cellular responses and drug delivery. We utilized *APOE3-* and *APOE4-* carrying BBB cells and investigated molecular changes following FUS^+MB^ as well as the FUS^+MB^-mediated delivery of two therapeutic AD antibodies: an Aducanumab analogue (anti-Aβ antibody, commercially known as Aduhelm^TM^) [24, 25], and RNF5, an anti-Tau antibody, which has recently been shown to be efficient in reducing p-Tau levels in an animal model with Tau pathology [26]. Here, we report the improved delivery of these antibodies in our BBB-like *in vitro* model using FUS^+MB^ and the cell-specific effects FUS^+MB^ confers in an *APOE3*/*APOE4* context.

## RESULTS

### Human iPSC-derived *APOE4* iBECs demonstrate key phenotypical differences compared to *APOE3* iBECs

To reliably model drug delivery in AD, we generated iBECs from previously published *APOE3* and *APOE4*-carrying human iPSCs, including one isogenic pair in which both *APOE4* alleles had been converted to *APOE3* using CRISPR-Cas9 [27-29]. iBECs were generated using a previously published protocol [18] and differentiation confirmed by the presence of BBB markers occludin, claudin-5 and ZO-1 in each line as well as the formation of characteristic cobblestone-like morphology (Fig. 1A and Fig. S1). To ensure physiologically relevant barrier integrity, TEER was measured in Transwells containing a membrane with pores of 0.4 μm in diameter (Ø) – a Transwell format most commonly used in iBEC-based models [18, 30, 31]. iBECs from all lines readily formed cobblestone-like monolayers on 0.4 μm pore Transwells, however *APOE4* iBECs demonstrated significantly reduced TEER (*P*<0.0001) compared to *APOE3* iBECs (1985±181 vs. 3056±66 Ohm/cm^2^, respectively, mean±SE; Fig. 1B). *APOE4* iBECs also exhibited higher variation in TEER measurements than *APOE3* lines, which clustered more closely together (Fig. 1B). To assess passive permeability of the formed barrier, leakage of fluorescently-conjugated 5 kDa dextran across the iBEC monolayer was measured, with significantly increased dextran permeability (*P*=0.0022) observed in *APOE4* iBECs compared to *APOE3* iBECs, supporting reduced barrier integrity of *APOE4* iBECs (Fig. 1B). To ensure differences in TEER and permeability were not caused by inefficient differentiation, we measured the relative gene expression of BBB markers occludin *(OCLN)*, claudin-5 *(CLDN5),* zonula occludens-1 *(TJP1),* VE-cadherin *(CDH5)* as well as the endothelial cell-specific SRY-box transcription factor 18 *(SOX18)* (Fig. 1C). Compared to undifferentiated iPSCs, all studied junctional markers were significantly upregulated (*P*<0.05) in *APOE3* and *APOE4* iBECs, with *SOX18* also being significantly increased (*P*=0.0096) in *APOE3* iBECs compared to iPSC lines and showing a trend towards an increase in *APOE4* iBECs (Fig. 1C). When the gene expression levels of the studied markers were compared between *APOE3* and *APOE4* iBECs, no significant differences were identified, suggesting a similar level of differentiation between the lines (Fig. 1D).

**Fig. 1.**
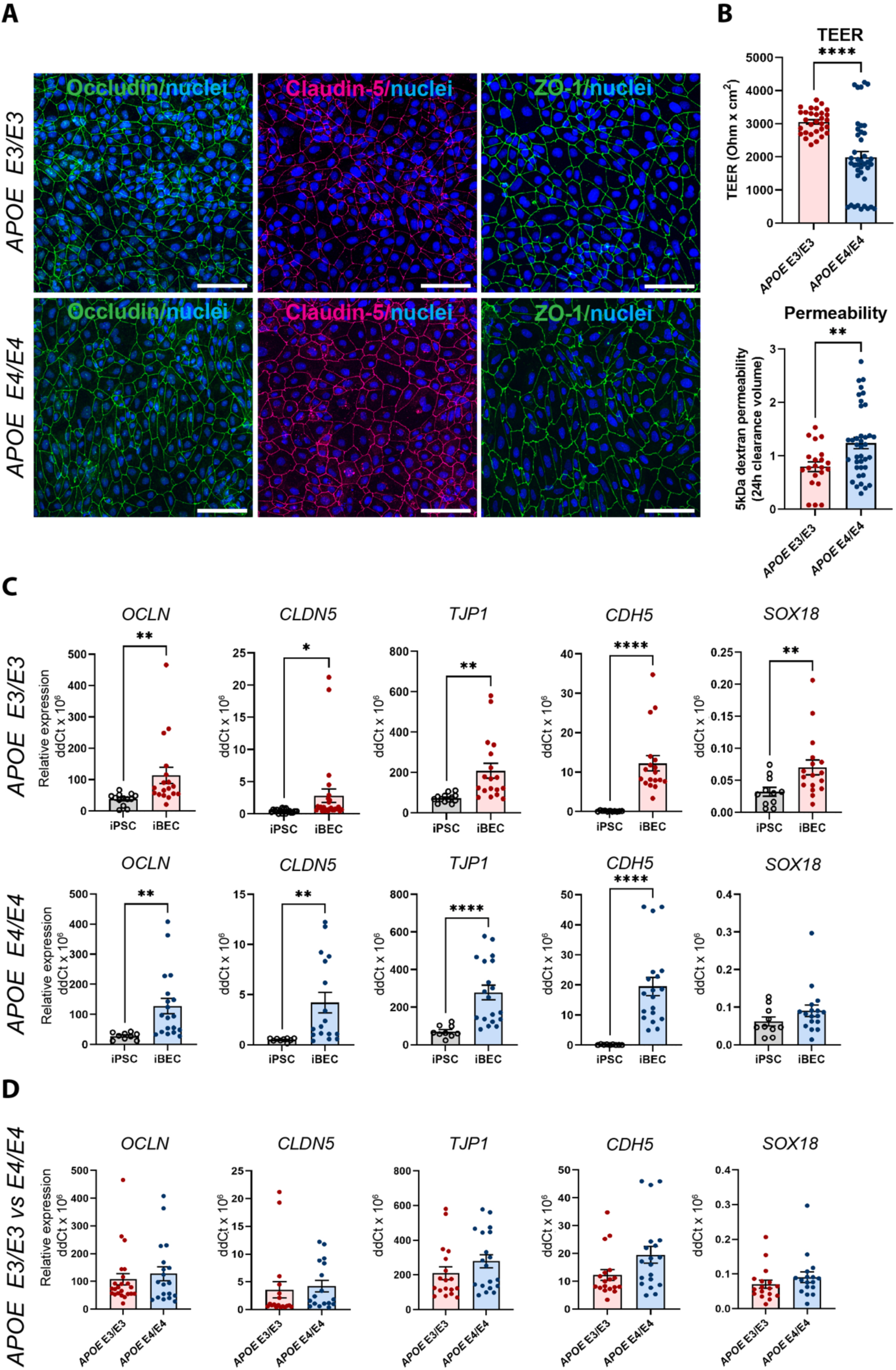
Characterization of brain endothelial-like cells (iBECs) from *APOE3* and *APOE4*-carrying human iPSCs. (**A**) Representative immunofluorescence images of Occludin (green), Claudin-5 (magenta) and ZO-1 (green) in *APOE3* and *APOE4* iPSC-derived iBECs (scale bar = 100 μm, nuclei stained using Hoechst). (**B**) Trans-endothelial electrical resistance (TEER, Ohm/cm^2^) and passive 5 kDa dextran permeability (24 h clearance volume) in *APOE3* and *APOE4* iBECs. (**C**) Relative gene expression of blood-brain barrier (BBB) and endothelial cell markers in *APOE3* iPSCs vs iBECs (top row) and *APOE4* iPSCs vs iBECs (bottom row). (**D**) Relative gene expression of BBB and endothelial cell markers in *APOE3* and *APOE4* iBECs. N=3 biological replicates and a minimum of n=3 independent replicates per line. \**P*<0.05, \*\**P*<0.01, \*\*\*\**P*<0.0001 by unpaired t-test with Welch’s correction, error bars = SEM.

### RNA sequencing reveals lack of transcriptome changes in *APOE3* and *APOE4* iBECs following FUS^+MB^

Previous animal studies have identified changes in the microvascular transcriptome following FUS^+MB^ [32]; however, molecular responses of the human BBB to FUS^+MB^ are largely unknown. Therefore, we performed bulk RNA-sequencing (RNA-seq) on FUS^+MB^-treated *APOE3* and *E4* iBECs. To achieve this, *APOE3* and *E4* iBECs were treated with FUS^+MB^ and collected at 1 h and 24 h post sonication, analogous to the previously reported timeline of BBB opening and spontaneous closing post FUS^+MB^ in the *in vitro* models [18, 33]. Corresponding untreated (UT) controls at the respective timepoints were included.

To gain insight into the relationships between the analyzed samples we first performed principal component analysis (PCA) on our RNA-seq dataset (Fig. 2A). This revealed that the largest variation between the samples was associated with *APOE3* and *APOE4* genotype (principal component 1, PC1) and sample collection time (PC2). Surprisingly, UT and FUS^+MB^ treated samples largely clustered together in donor pairs, providing the first indication that FUS^+MB^ had little effect on the iBEC transcriptome. Indeed, differential expression analysis identified no differentially expressed genes (DEGs) between FUS^+MB^ and UT at either timepoint analyzed irrespective of the *APOE* genotype (Fig. 2B), suggesting that the effects of FUS^+MB^ on gene expression in human iBECs are minimal. Gene expression also did not differ when *APOE* genotype was incorporated into our analysis (Fig. S2), indicating that the presence of *APOE E3* or *E4* allele does not have a profound effect on iBEC responses to FUS^+MB^. We then compared UT *APOE3* and UT *APOEE4* iBECs at 1 h which interestingly revealed 635 DEGs (FDR < 0.05), with 405 significantly up-regulated and 230 significantly down-regulated genes in *APOE4* iBECs compared to *APOE3* iBECs (Fig. 2C, Table S1). Intriguingly, several genes identified as the most up- or down-regulated in *APOE4* iBECs, including *SHMT1* [34], *HMGB2* [35], *ALDH7A1* [36], *ABL2* [37], have previously been linked to cognitive dysfunction and AD, although their role at the BBB requires further investigation. Finally, to explore the biological function of identified DEGs between UT *APOE4* and *APOE3* iBECs at 1 h, we conducted gene ontology (GO) enrichment analysis for biological processes. The results revealed that the DEGs were significantly enriched in 302 GO terms (Table S2) with the top three enriched terms being “DNA replication”, “DNA-dependent DNA replication” and “chromosome segregation” (Fig. 2D, Fig. S3) suggesting alterations in DNA and cell division processes in *APOE4* iBECs.

**Fig. 2.**
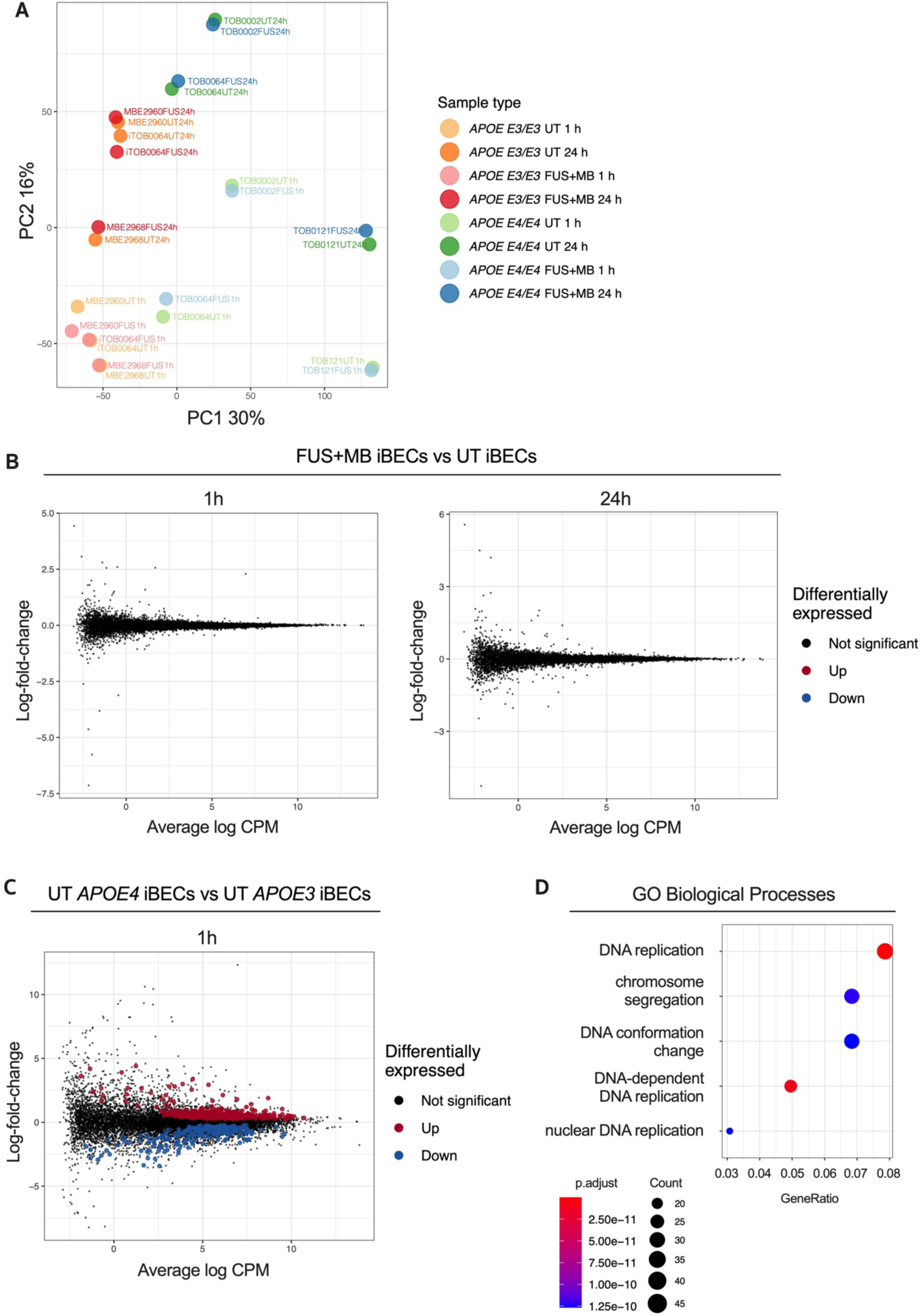
Transcriptome analysis of the response to FUS^+MB^ in *APOE3* and *APOE4* iBECs. (**A**) Principal component analysis (PCA) of gene expression profiles for all iBEC samples. The percentage of variance explained by principal component 1 (PC1) and principal component 2 (PC2) shown in axis labels. (**B**) Mean-difference (MD) plots showing log-fold-change vs average log expression values (log2 counts per million, CPM). Left panel: FUS^+MB^ treated *APOE3* and *APOE4* iBECs at 1 h vs UT *APOE3* and *APOE4* iBECs at 1 h. Right panel: FUS^+MB^ treated *APOE3* and *APOE4* iBECs at 24 h vs UT *APOE3* and *APOE4* iBECs at 24 h. (**C**) MD plot showing log-fold-change and average log expression values in UT *APOE4* iBECs at 1 h versus *APOE3* iBECs at 1 h. N=3 biological replicates. (**D**) Dot plot of top 5 gene ontology (GO) terms from sub ontology Biological Process enriched from comparison of UT *APOE4* iBECs at 1h vs UT *APOE3* iBECs at 1 h. The dot size represents the number of genes associated with the GO term and the dot color represents the FDR value. The differentially expressed genes (i.e. FDR <0.05) from this comparison were used as input to the analysis.

### FUS^+MB^ increases the *in vitro* delivery of potentially therapeutic Alzheimer’s antibodies Aducanumab and anti-Tau (RNF5)

Although the classically used Ø 0.4 µm pore Transwells when coated with collagen IV and fibronectin (in the absence of iBEC) readily allow the passage of small molecule (5 kDa) dextran, our preliminary experiments revealed that this format of inserts drastically limits the permeability of the large molecule (150 kDa) dextran, which corresponds to therapeutic antibody size. To enable increased antibody delivery through the Transwell membrane itself, we established a new Transwell model whereby iBECs were cultured on wider, Ø 3.0 µm pore Transwell inserts with the use of these inserts alone (containing collagen IV and fibronectin coating but no iBECs) resulting in a 7.5 fold increase in 150 kDa dextran transfer through the membrane compared to 0.4 μm inserts (Fig. S4A). By further increasing the seeding cell number by 1.5 fold, compared to cell number used for the 0.4 μm pore Transwells, we were also able to generate classical ‘cobblestone’ forming iBEC monolayers in 3.0 μm pore inserts assessed by ZO-1 localization to TJs (Fig. 3A and Fig. S4B). In addition, we were able to obtain TEER values for *APOE3* (1609±46 Ohm/cm^2^, mean±SE) and *APOE4* iBECs (672±29 Ohm/cm^2^, mean±SE), which albeit lower than in 0.4 μm pore Transwells, were still substantially higher than those reported for any primary or immortalized BEC-based Transwell model [38]. Consistent with the findings in 0.4 μm pore Transwells reported above, TEER values in 3.0 μm pore Transwells were significantly lower (*P*<0.0001) for *APOE4* iBECs compared to *APOE3* iBECs with *APOE4* iBECs showing a trend towards increased passive permeability to 150 kDa dextran (Fig. 3B).

**Fig. 3.**
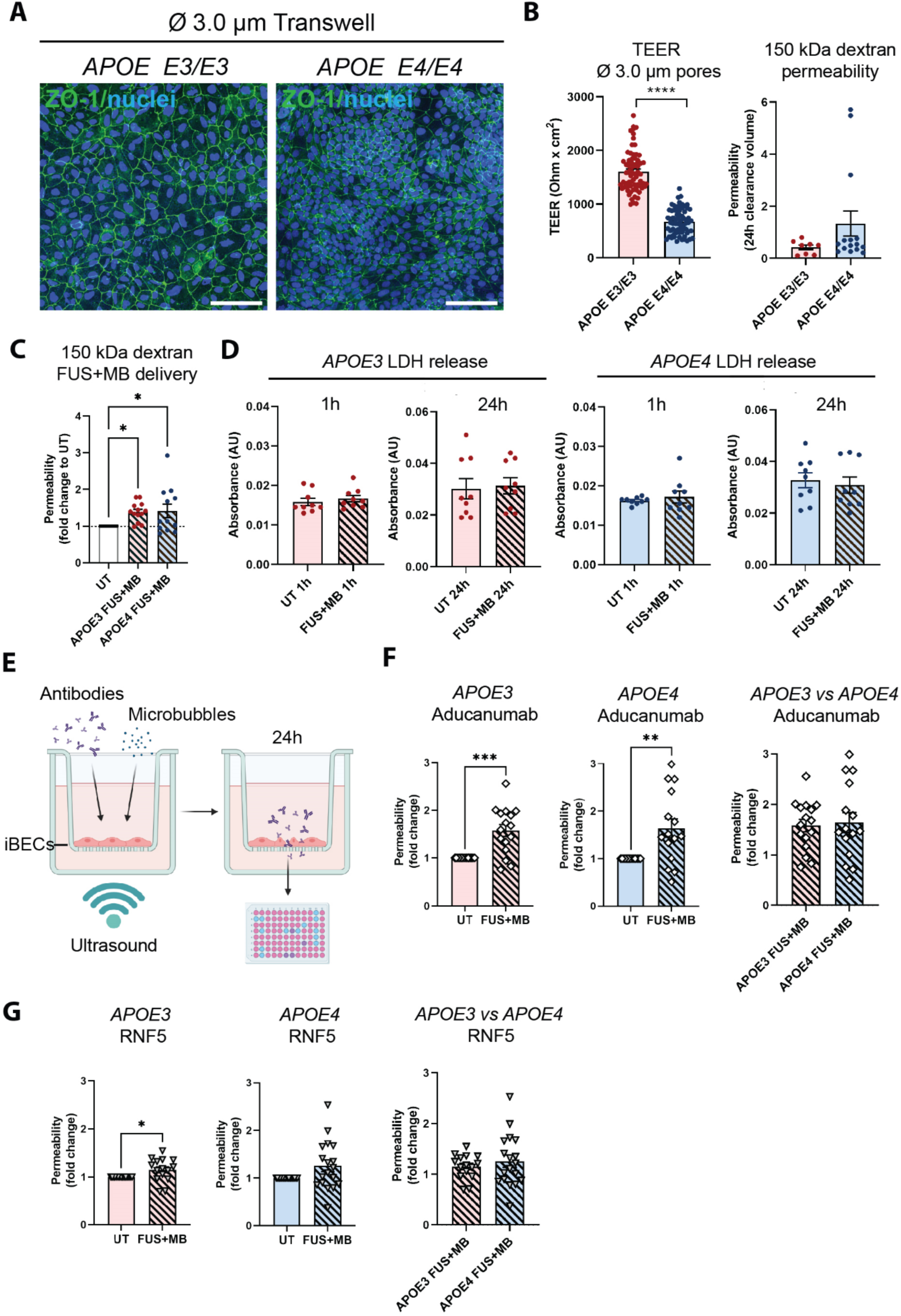
Therapeutic antibody delivery in *APOE3* and *APOE4* iBEC monocultures following FUS^+MB^. (**A**) Immunofluorescence images of iBECs cultured in Ø 3.0 μm Transwell stained with ZO-1 (green, Hoechst counterstain, scale bar = 100 μm). (**B**) Trans-endothelial electrical resistance (TEER, Ohm/cm^2^) and passive 150 kDa dextran permeability (24 h clearance volume) in *APOE3* and *APOE4* iBECs in 3.0 μm Transwells. (**C**) Delivery of 150 kDa dextran in *APOE3* and *APOE4* iBECs using focused ultrasound + microbubbles (FUS^+MB^). Permeability shown as fold change to untreated (UT) at 24h. (**D**) Lactate dehydrogenase (LDH) release (shown as absorbance, AU) in UT and FUS^+MB^ treated *APOE3* and *APOE4* iBECs at 1 h and 24 h following treatment. (**E**) Schematic illustration of therapeutic antibody delivery using FUS^+MB^ in the Transwell system. Fluorescently-conjugated antibodies were added together with MBs to a Transwell containing iBECs and the insert was exposed to ultrasound. Flow-through of antibodies was measured 24 h following the treatment using a fluorescent plate reader. Graphic created using Biorender.com. (**F**) Aducanumab delivery in UT and FUS^+MB^ treated *APOE3* and *APOE4* iBECs as well as comparison of Aducanumab delivery in FUS^+MB^ treated *APOE3* vs *APOE4* iBECs (permeability shown as fold change to UT at 24 h). (**G**) RNF5 delivery in UT and FUS^+MB^ treated *APOE3* and *APOE4* iBECs, as well as comparison of RNF5 delivery in FUS^+MB^ treated *APOE3* vs *APOE4* iBECs. Permeability shown as fold change to UT at 24 h. N=3 biological replicates and minimum n=3 independent replicates per cell line. \**P*<0.05, \*\**P*<0.01, \*\*\**P*<0.001, \*\*\*\**P*<0.0001 by unpaired t-test with Welch’s correction, error bars = SEM.

Using previously established FUS^+MB^ parameters [18], we next assessed 150 kDa dextran permeability following exposure to FUS^+MB^. When FUS^+MB^ was applied to *APOE3* and *APOE4* iBEC monolayers cultured on 3.0 μm pore Transwells, this demonstrated significantly increased delivery of 150 kDa dextran (*P*<0.05) compared to untreated (UT) cells (Fig. 3C). To assess the effects of FUS^+MB^ on cell viability we used a lactate dehydrogenase (LDH) cytotoxicity assay; however, we did not detect differences in LDH levels between samples collected from UT and FUS^+MB^ treated cells in *APOE3* or *APOE4* iBECs, suggesting FUS^+MB^ do not affect cell viability (Fig. 3D).

Following optimization of the 3.0 μm Transwell platform for FUS^+MB^-mediated drug delivery, we proceeded to measure the delivery of fluorescently-conjugated Aducanumab (anti-amyloid-β) and RNF5 (anti-Tau) therapeutic antibodies in our model system. Briefly, 48 h following purification on 3.0 μm Transwells, iBECs were simultaneously treated with 1μM (150 μg/ml) of the antibody of interest along with MBs and exposed to FUS at 0.3 MPa (286 kHz centre frequency) for 120 s (Fig. 3E). Fluorescence of the flow-through, indicating antibody delivery, was measured 24 h following treatment (Fig. 3E). Interestingly, Aducanumab delivery was significantly increased in both *APOE3* (1.58±0.12 fold, mean±SE, *P*=0.0003) and *APOE4* (1.64±0.2 fold, mean±SE, *P*=0.0059) iBEC cultures following FUS^+MB^ treatment compared to UT cells (Fig. 3F). The delivery efficiency of Aducanumab following FUS^+MB^ was not significantly different between *APOE3* and *APOE4* iBECs. Interestingly, the delivery of RNF5 was significantly increased in *APOE3* (1.15±0.06 fold, mean±SE, *P*=0.0295) with *APOE4* iBECs demonstrating a strong trend towards increase in RNF5 delivery (1.26±0.12 fold, mean±SE, *P*=0.0501) when treated with FUS^+MB^ as compared to UT (Fig. 3G). Similar to Aducanumab, RNF5 delivery in FUS^+MB^ treated *APOE3* and *APOE4* iBECs did not significantly differ (Fig. 3G).

### FUS^+MB^ treatment has minimal effects on *APOE3* and *APOE4* iAstrocyte phenotype

Astrocytes are a critical component of the BBB, but little is known about the effects of FUS^+MB^ on human astrocytes. We therefore examined astrocyte responses to FUS^+MB^. We generated induced astrocytes (iAstrocytes) from the same *APOE3* and *APOE4*-carrying human iPSCs that were used for iBEC differentiation (n=2 biological replicates including 1 isogenic pair). Neural progenitor cells (NPCs) were generated and characterized for nestin and SOX2 expression (Fig. S5A) and then differentiated into iAstrocytes for a minimum of 60 days. iAstrocytes were matured 7 days before experiments with 10 ng/mL bone morphogenetic protein 4 (BMP-4) and ciliary neurotrophic factor (CNTF) as previously described [39]. Matured iAstrocytes were exposed to FUS^+MB^ using the same parameters as for therapeutic antibody delivery in iBEC monocultures, and cell morphology, viability, marker expression and inflammatory responses were examined 1 h and 24 h following treatment.

Following FUS^+MB^ treatment, *APOE3* and *APOE4* iAstrocytes did not display any adverse morphological changes compared to UT cells when stained with astrocyte markers aquaporin-4 (AQP4) and glial fibrillary acidic protein (GFAP) 1 h and 24 h following treatment (Fig. 4A and Fig. S5B). In addition, the LDH assay did not reveal any significant changes in cell viability between UT and FUS^+MB^ conditions at 1 h and 24 h timepoints in *APOE3* or *APOE4* iAstrocytes (Fig. 4B). Next, we analyzed the relative gene expression of astrocyte markers following FUS^+MB^ treatment. No significant changes in expression of astrocyte markers *AQP4*, *GFAP* (Fig. 4C) and *S100B* (Fig. S6A) were identified in FUS^+MB^ treated *APOE3* or *APOE4* iAstrocytes compared to UT iAstrocytes. Following this, to identify whether FUS^+MB^ elicited any inflammatory responses in iAstrocytes, we examined the gene expression of inflammatory cytokines known to be secreted by iPSC-derived astrocytes in basal culture conditions and following inflammatory stimulation [40-42]. Interestingly, whereas interleukin-1β (*IL-1β*) expression was not altered in *APOE3* or *APOE4* iAstrocytes 1h following FUS^+MB^ treatment when compared to UT, its expression was significantly reduced (*P*=0.0085) in *APOE3* iAstrocytes at 24 h following FUS^+MB^ treatment (Fig. 4D). Similarly, *IL-6* and *IL-8* expression was not altered in *APOE3* or *APOE4* iAstrocytes 1 h following FUS^+MB^, however, their expression levels were significantly (*IL-6*: *P*=0.0447; *IL-8 P*=0.0077) downregulated in *APOE3* iAstrocytes 24 h following FUS^+MB^ treatment, remaining unchanged in *APOE4* iAstrocytes following FUS^+MB^ treatment (Fig. 4D). No changes were identified in the expression of C-C motif chemokine ligand 2 (*CCL2*), an astrocytic chemokine acting as an important mediator of damage-associated neuroinflammation [43, 44], in *APOE3* or *APOE4* following FUS^+MB^ treatment for either timepoint (Fig. S6B). Finally, we compared cytokine expression in a FUS^+MB^-treated isogenic *APOE3*/*APOE4* iAstrocyte pair (parent *APOE4* iPSC line converted to *APOE3* [27]). Interestingly, the results demonstrated an increased inflammatory response in *APOE4* iAstrocytes compared to *iAPOE3* cells 1 h after treatment, with significantly increased *IL-1β* (*P*=0.025), *IL-8* (*P*=0.017) and *CCL2* (*P*=0.043) expression observed in *APOE4* iAstrocytes (Fig. S6C). At 24 h, these differences were no longer evident, but with *IL-8* expression significantly decreased (*P*=0.047) in *APOE4* vs *iAPOE3* iAstrocytes.

**Fig. 4.**
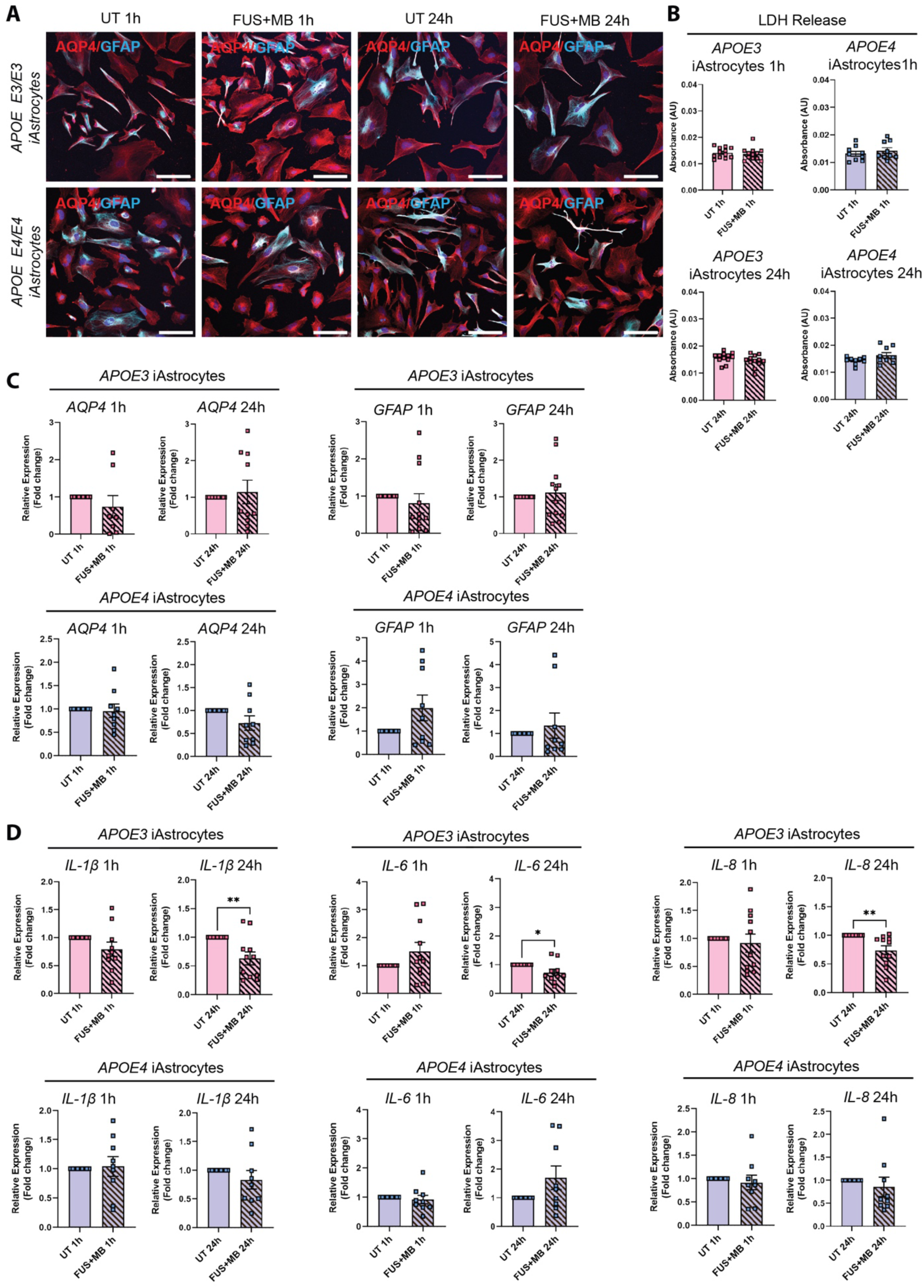
Effect of focused ultrasound on *APOE3* and *APOE4* iAstrocytes. (**A**) Representative immunofluorescence images of *APOE3* and *APOE4* iAstrocytes in untreated (UT) and focused ultrasound + microbubble (FUS^+MB^) conditions 1 h and 24 h following treatment stained with AQP4 (red) and GFAP (cyan; Hoechst counterstain, scale bar = 100 μm). (**B**) Lactate dehydrogenase (LDH) release (shown as absorbance, AU) in UT and FUS^+MB^ treated *APOE3* and *APOE4* iAstrocytes 1 h and 24 h following treatment. (**C**) Relative gene expression of astrocyte markers *AQP4* and *GFAP* in UT and FUS^+MB^ treated *APOE3* and *APOE4* iAstrocytes 1 h and 24 h after treatment. (**D**) Relative gene expression of inflammatory markers *IL-1β*, *IL-8* and *IL-6* in UT and FUS^+MB^ treated *APOE3* and *APOE4* iAstrocytes 1 h and 24 h after treatment. N=2 biological replicates and minimum n=3 independent replicates per line. \**P*<0.05, \*\**P*<0.01 by unpaired t-test with Welch’s correction, error bars = SEM.

### iBECs co-cultured with iAstrocytes demonstrate increased barrier integrity and allow therapeutic antibody delivery following FUS^+MB^

As a step towards developing a multicellular AD BBB model for FUS^+MB^ mediated therapeutic antibody delivery, we next established iBEC+iAstrocyte co-cultures in 3.0 μm pore Transwells as astrocytes are known to be key players in enhancing BBB integrity [45]. Cells were co-cultured for 24 h prior to performing antibody delivery experiments, and with sample collection performed 24 h following treatment the total time in co-culture was 48 h. Both cell types were maintained in endothelial serum-free medium (ESFM) + B+27 supplement, with the co-culture conditions supporting normal marker expression of both cell types (Fig. 5A). Twenty-four hours following co-culture, TEER was significantly increased in both *APOE3* (2147±145 Ohm/cm^2^, mean±SE, *P*=0.0082) and *APOE4* (1504±80 Ohm/cm^2^, mean±SE, *P*<0.0001) co-cultures when compared to the respective iBEC monocultures (*APOE3* iBECs: 1609±46 Ohm/cm2, mean±SE; *APOE4* iBECs: 672±29 Ohm/cm2, mean±SE; Fig 4B) in 3.0 μm pore Transwells, supporting barrier enhancing effects of iAstrocytes (Fig. 5B). Interestingly, when we compared TEER between *APOE3* and *APOE4* co-cultures, TEER was significantly reduced (*P*=0.0004) in *APOE4* co-cultures compared to *APOE3* co-cultures, in line with our findings from iBEC monoculture experiments. Next, we performed therapeutic AD antibody delivery in iBEC+iAstrocyte co-cultures with FUS^+MB^ and measured delivery efficiency 24 h following treatment (Fig. 5C). Similar to iBEC monocultures, Aducanumab delivery was significantly increased in both *APOE3* (1.73±0.39 fold, mean±SE, *P*=0.0005) and *APOE4* (1.29±0.1, mean±SE, *P*=0.012) co-cultures (Fig. 5D). Interestingly, the delivery of Aducanumab following FUS^+MB^ was significantly lower (*P*=0.017) in *APOE4* co-cultures compared to *APOE3* co-cultures (Fig. 5D). When compared to iBEC monocultures, there was no difference in Aducanumab delivery efficiency in *APOE3* co-cultures, while in contrast, *APOE4* co-cultures demonstrated significantly lower (*P*=0.012) Aducanumab delivery when compared to *APOE4* monocultures (Fig. 5E). When RNF5 delivery following FUS^+MB^ was analyzed in *APOE3* and *APOE4* co-cultures, it was significantly increased in both culture types compared to UT (*APOE3*: 1.50±0.2 fold, *P*=0.038; *APOE4* 1.55±0.09 fold, *P*<0.0001, mean±SE) and no significant difference in RNF5 delivery between *APOE3* and *APOE4* co-cultures was observed (Fig. 5F). Consistent with Aducanumab, the delivery efficiency of RNF5 was similar between *APOE3* co- and iBEC monocultures following FUS^+MB^ (Fig. 5G). In contrast, compared to iBEC monocultures, RNF5 delivery efficiency following FUS^+MB^ was significantly increased (*P*=0.018) in *APOE4* co-cultures, suggesting possible modifying effects of iAstrocytes (Fig. 5G).

**Fig. 5.**
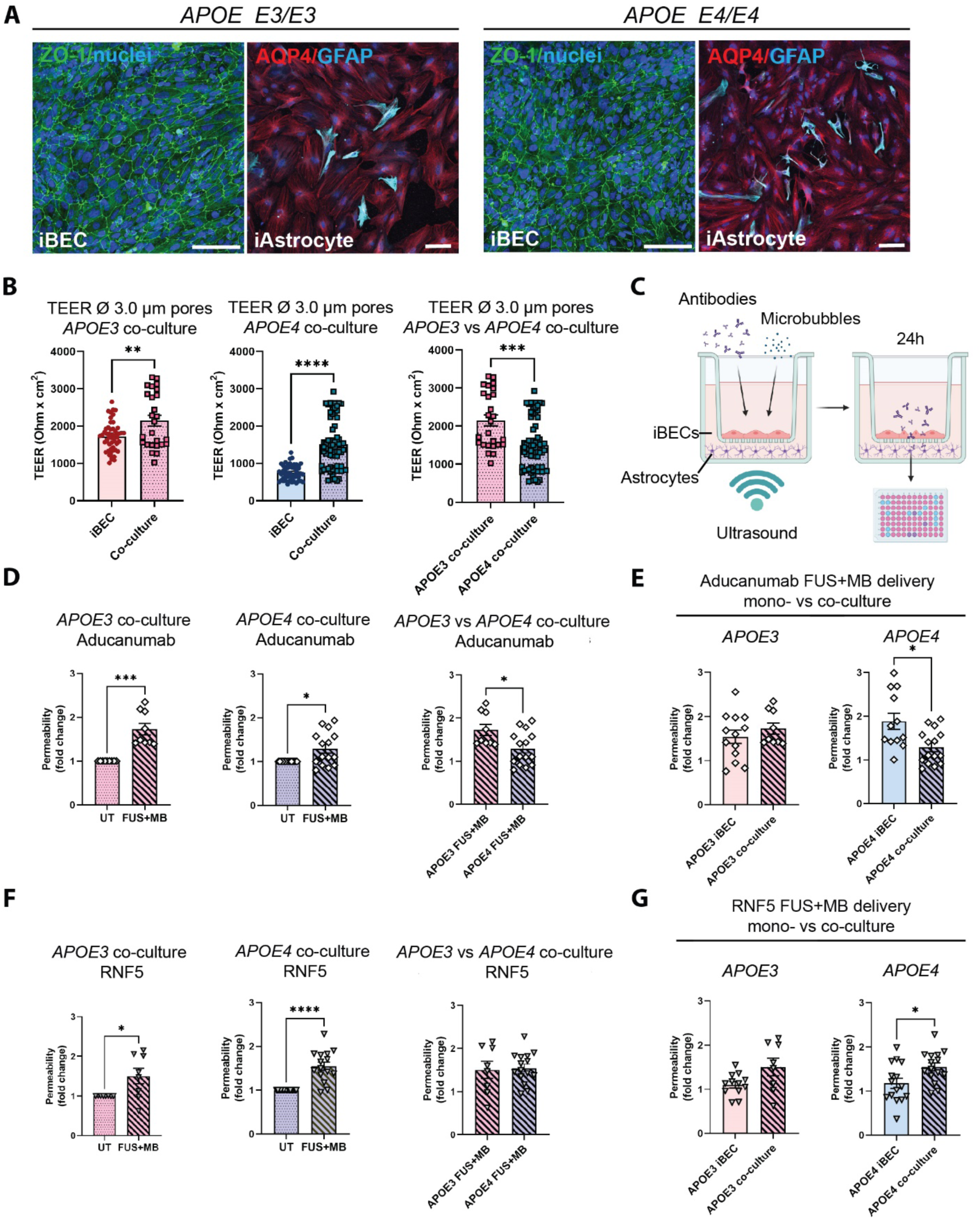
Therapeutic AD antibody delivery in *APOE3* and *APOE4* iBEC+iAstrocyte co-cultures following FUS^+MB^. (**A**) Representative immunofluorescence images of *APOE3* and *APOE4* iBECs (ZO-1/green) and iAstrocytes (AQP4/red, GFAP/cyan) in co-culture, maintained in endothelial serum free medium + B-27 (Hoechst counterstain, scale bar = 100 μm). (**B**) Comparison of trans-endothelial electrical resistance (TEER, Ohm/cm^2^) in *APOE3* and *APOE4* iBECs and iBEC+astrocyte co-cultures as well as *APOE3* vs *APOE4* co-cultures in Ø 3.0 μm Transwells. (**C**) Schematic illustration of therapeutic antibody delivery using FUS^+MB^ in iBEC+iAstrocyte co-cultures. iBECs+iAstrocytes were co-cultured in the Transwell system for 24 h after which antibodies were added together with MBs and insert exposed to ultrasound. Flow-through of antibodies was measured 24h following treatment using a fluorescence plate reader. Graphic created using Biorender.com. (**D**) Aducanumab delivery in UT and FUS^+MB^ treated *APOE3* and *APOE4* iBEC+iAstrocyte co-cultures, as well as comparison of Aducanumab delivery in FUS^+MB^ treated *APOE3* vs *APOE4* iBEC+iAstrocyte co-cultures. Permeability shown as fold change to UT at 24 h. (**E**) Comparison of Aducanumab delivery following FUS^+MB^ in *APOE3* and *APOE4* iBEC mono- and iBEC+iAstrocyte co-cultures. (**F**) RNF5 delivery in UT and FUS^+MB^ treated *APOE3* and *APOE4* iBEC+iAstrocyte co-cultures, as well as comparison of RNF5 delivery in FUS^+MB^ treated *APOE3* vs *APOE4* iBEC+iAstrocyte co-cultures. Permeability shown as fold change to UT at 24 h. (**G**) Comparison of RNF5 delivery following FUS^+MB^ in *APOE3* and *APOE4* iBEC mono- and iBEC+iAstrocyte co-cultures. N=2 biological replicates and minimum n=3 independent replicates per cell line. \**P*<0.05, \*\*\**P*<0.001, \*\*\*\**P*<0.0001 by unpaired t-test with Welch’s correction, error bars = SEM.

### A novel hydrogel-based 2.5D BBB model provides a physiologically relevant alternative to the Transwell system for the high-throughput drug delivery studies

While the Transwell system for drug permeability screening is easy to establish and widely used, it requires large volumes of reagents (such as culture medium and antibodies), and cells in co-culture are separated by an artificial membrane. Additionally, the inherent 6- to 24-well format of the Transwell system limits its use for high-throughput drug screening. To overcome these issues, we developed a 2.5D gel-based BBB-like model in a 96-well plate that could offer a more physiologically relevant alternative where astrocytes are grown in 3D and iBECs in 2D, and are in contact with each other without a separating membrane (Fig. 6A). In addition, the reagent requirements for high-throughput screening of antibodies or alternative drug delivery methods are downscaled in this model. Matrigel and collagen I are commonly used for 3D assays, however, they often display batch-to-batch variability and are too soft for establishing a BBB model. Thus, we trialed LunaGel^TM^, a gelatin-based photocrosslinkable gel, which can be adjusted to a desired stiffness using varying polymerization times under blue light (Gelomics). Briefly, to establish a 3D culture we first embedded iAstrocytes in LunaGel^TM^ and matured iAstrocytes for 7 days. Next, iBECs were purified on collagen IV and fibronectin yielding a highly homogeneous BEC-like population as shown by us and others [18, 30]). The iBECs were subsequently seeded in 2D on top of the iAstrocyte layer, on a new layer of high-stiffness LunaGel^TM^ and allowed to attach for 24 h. The 2.5D co-culture model was then exposed to antibodies, MBs and FUS as per the Transwell system and fluorescence intensity was measured in the gel 24 h following treatment using a plate reader (Fig. 6A). Interestingly, our results demonstrated that when using low stiffness LunaGel^TM^, iAstrocytes readily proliferated and extended their processes within the gel (Fig. 6B). Following addition of a thin layer of high stiffness LunaGel^TM^ on top of the iAstrocyte gel layer and coating with collagen IV and fibronectin, purified iBECs attached on the iAstrocyte layer in medium supplemented with factors known to promote endothelial cell adhesion [46, 47], thereby forming a uniform barrier (Fig. 6B). Intriguingly, our initial observations have demonstrated significantly increased (*P*=0.033) Aducanumab delivery following FUS^+MB^ (fold change 1.4±0.12, mean±SE), when compared to UT (Fig. 6C). Although further optimization of this model is on-going, our results highlight the potential of using a gel-based 2.5D model system for FUS^+MB^-mediated antibody delivery as a more physiologically relevant high-throughput platform with potentially higher translatability to patients than with the traditional Transwell system.

**Fig. 6.**
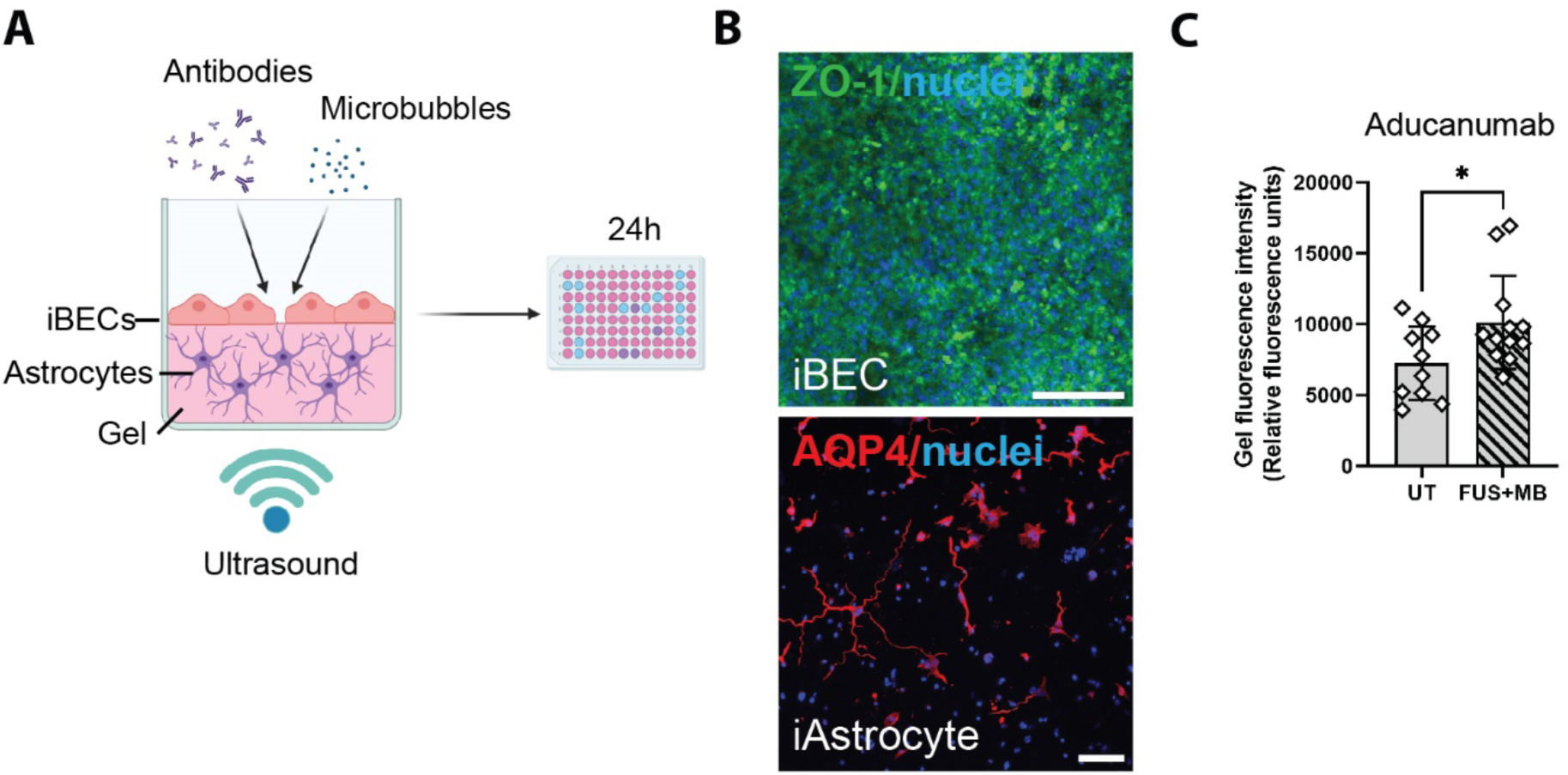
A 2.5D gel-based BBB model allows for screening of FUS^+MB^-mediated antibody delivery. (**A**) A schematic illustration of therapeutic antibody delivery using FUS^+MB^ in LunaGel^TM^-based 2.5D model. iAstrocytes were embedded in the gel and allowed to mature for 7 days. Following purification yielding a highly homogeneous population of iBECs, the cells were seeded on top of the iAstrocyte layer and exposed to therapeutic antibodies, MB and ultrasound. Fluorescence intensity within the gel, indicating antibody delivery, was measured 24 h following treatment using a fluorescence plate reader. Graphic created using Biorender.com. (**B**) Representative immunofluorescence images of iBECs (ZO1/green) and iAstrocytes in the LunaGel^TM^ platform. Hoechst counterstain, scale bar = 100 μm. (**C**) Aducanumab delivery in UT and FUS^+MB^ treated iBEC+iAstrocyte 2.5D model indicated as relative fluorescence units. N=3 biological replicates. \**P*<0.05, by unpaired t-test with Welch’s correction, error bars = SEM.

## DISCUSSION

A major hindrance in treating neurodegenerative diseases such as AD is the low bioavailability of therapeutics in the brain due to the BBB, which inhibits the entry of most large molecule drugs, including antibodies, into the CNS. The FUS^+MB^ technique is a relatively novel technology that allows the transient opening of the BBB to improve the delivery of drugs into the CNS. A small number of clinical studies using FUS^+MB^ in AD patients have demonstrated the safety of this technique, making it a promising tool for increased drug delivery [20, 21, 48, 49]. However, our understanding of the molecular effects of FUS^+MB^ on human BBB cells is limited, and there is a lack of patient cell-based screening platforms to identify FUS-deliverable drugs. Thus, patient cell models that recapitulate disease phenotype and allow for the testing of FUS^+MB^ effects are critical in accelerating the translation of FUS^+MB^ to the clinic for the treatment of a range of neurodegenerative diseases.

We have previously demonstrated the ability to increase BBB permeability and Aβ clearance using FUS^+MB^ in a patient cell Transwell-based model that incorporated iBECs carrying the familial AD *PSEN1* mutation [18]. However, with familial AD accounting for less than 5% of all AD cases, this model is not highly representative of the majority of AD cases which are of sporadic origin. Thus, with the known association of the sporadic AD risk gene polymorphism *APOE4* on BBB dysfunction, in this study we utilized *APOE4*-carrying patient-derived iPSCs to generate a BBB-like model to test the effects of FUS^+MB^ as well as the delivery of two therapeutic AD antibodies and compared the results to *APOE3* (normal AD risk control) cells. Our hypothesis was that a patient-specific sporadic AD cell model of the BBB will increase our understanding of FUS^+MB^ effects that are applicable to the majority of AD cases, thereby accelerating the translation of FUS^+MB^ into a clinical drug delivery method for the treatment of AD.

Homozygote carriers of the *APOE4* allele have 15-fold increased risk of developing AD compared to the most common isoform *APOE3* when homozygous [2]. Both animal and human studies have reported an association between *APOE4* and a dysfunctional BBB [8, 11]. This is important to consider as a dysfunctional BBB alters the physiological conditions in the brain and can further hinder drug delivery by changing how the drug is taken up by diffusion or metabolized in the brain [6]. Thus, it is vital to understand the contribution of a disease polymorphism on drug delivery efficiency. Using patient-derived iPSCs is important as they can complement findings from transgenic mouse models, which can lack human disease resemblance and demonstrate structural and molecular differences to the human BBB [50-53]. Although questions have been raised about the accuracy of iPSC-derived iBECs in resembling *in vivo* BECs [54], these cells exhibit the highest TEER compared to any other BEC model making them ideal for mechanistic BBB opening studies such as FUS^+MB^ [16, 38]. In addition, iBECs exhibit many properties of endogenous BECs and most closely resemble the permeability of the human BBB when compared to other cell models [16, 31], further supporting their use. As such, we hypothesized that iBECs are the most appropriate cell model for FUS^+MB^-mediated drug delivery modelling *in vitro* in an AD-context and opted for their use in this study.

Our characterization of *APOE3* and *APOE4* iBECs revealed several phenotypical differences that indicate the contribution of *APOE4* to BBB breakdown. Although both *APOE3* and *APOE4* iBECs generated a cobblestone-like morphology and expressed key BBB markers, the TEER of *APOE4* iBECs was significantly lower compared to *APOE3* iBECs in both mono- and iAstrocyte co-culture systems, supporting the association of *APOE4* with reduced BBB integrity. Reduced TEER observed in *APOE4* iBECs also correlated with increased permeability to a small molecule fluorescent tracer, in-line with the reported association of *APOE4* with BBB breakdown [11]. Overall, our results demonstrate the ability to model *APOE4*-induced BBB changes in an *in vitro* patient-cell model. Importantly, our results did not identify gene expression differences in BBB-associated TJs and AJs between *APOE3* and *APOE4* iBECs, suggesting the differences observed in BBB integrity between *APOE3* and *APOE4* cells are not driven by TJP or AJ expression, but rather alterations in TJP and AJ molecular architecture.

As the molecular effects of FUS^+MB^ on human BBB cells are unknown, we investigated the iBEC transcriptome following FUS^+MB^ exposure. Intriguingly, we did not identify any genes that were significantly altered by FUS^+MB^ irrespective of disease phenotype. In line with our study, using single-cell RNA sequencing, Gorick and colleagues also detected only minimal changes (8 DEGs overall) in the transcriptome of the murine cerebrovascular endothelium when FUS^+MB^ was applied at low pressure (0.1, 0.2 and 0.4 MPa) [55]. This could indicate FUS^+MB^ achieves the BBB opening by affecting junctional protein ultrastructure in BECs as previously suggested [56, 57], rather than by acting at the TJ or AJ gene expression level. Interestingly, a previous study conducted in a murine model identified changes in the microvessel transcriptome following exposure to FUS^+MB^, pointing to a transient inflammatory response induced by sonication [32]. Importantly, in contrast to our study, McMahon *et al*. [32] performed their analysis on dissected brain microvessels suggesting that the observed effects can be driven by other cells associated with microvasculature rather than just BECs. Correspondingly, our study also identified significant changes in gene expression levels of inflammatory mediators *IL-1β*, *IL-6* and *IL-8* in iAstrocytes following sonication, and Leinenga *et al.* has recently reported changes in the microglial transcriptome following FUS^+MB^ [58]. Lack of observed effects in our study could also be explained by differences in FUS parameters and MB type/concentration applied as compared to McMahon *et al*. and Gorick *et al*., illustrating the importance of further understanding the correlation between FUS and MB physical characteristics and cell responses. Finally, interspecies differences reported in transcriptome of murine and human microvessels [52], as well as high variation between patient-derived iPSC lines [27], could underlie the lack of identified changes to the transcriptome following FUS^+MB^. Overall, as we did not identify any adverse effects of FUS^+MB^ on iBEC viability nor effects on the iBEC transcriptome, our study further supports the safety of FUS^+MB^ treatment.

Interestingly, our analysis identified over 600 DEGs in *APOE3* vs *APOE4* iBECs, and further analysis of these genes may potentially identify novel disease biomarkers. Correspondingly, our GO enrichment analysis suggests DNA replication and cell division changes in the *APOE4* iBECs. Intriguingly, defects in mitosis and chromosome segregation have been previously linked to AD pathogenesis, with *APOE4* being one of the suggested genetic drivers of this effect [59-63]. Although alterations in BEC division have not yet been investigated in the context of BBB breakdown in AD, and only low level of cell turnover has been previously reported in iBECs and BECs *in vivo* under normal conditions [64, 65], our observation might point to a novel pathomechanism driving BBB dysfunction in *APOE4* carriers.

Large molecule therapeutic antibodies are a promising treatment for various neurodegenerative diseases, but their size limits their effective delivery across the BBB into the brain. It is estimated that only 0.2% of the intravenously administered antibody concentration reaches the brain, severely limiting their therapeutic efficacy [66]. As such, due to the BBB opening effects of FUS^+MB^ [67, 68], this technique provides a highly promising approach for large molecule therapeutic antibody delivery into the brain [69-71]. With there being a lack of *in vitro* cell models for FUS^+MB^-mediated drug permeability screening, we tested our established *APOE3* and *APOE4* iBEC platform for FUS^+MB^-mediated AD antibody delivery using a similar approach to our previously published study [18]. Interestingly, our results demonstrated a significant fold increase in Aducanumab and RNF5 antibody delivery in both *APOE3* and *APOE4* iBEC mono- and co-culture models. This first ever patient-derived cell-based model for FUS^+MB^-mediated therapeutic AD antibody delivery provides evidence of the promising effects of FUS^+MB^ in therapeutic antibody delivery. In clinical trials, Aducanumab has been shown to significantly inhibit cognitive decline when administered at the highest possible dose [72]. Thus, increasing the concentration of Aducanumab reaching the brain with methods like FUS^+MB^ could further improve clinical outcomes.

Astrocytes are central components of the BBB, enhancing barrier integrity and providing communication with the BBB and CNS [45]. In addition, astrocytes are key mediators of inflammation by secreting cytokines that in turn modulate other brain cells, such as microglia [73]. The effects of FUS^+MB^ on human astrocytes have been sparsely investigated, thus we wanted to examine potential modulatory effects of FUS^+MB^ on iAstrocytes and whether *APOE3* and *APOE4* iAstrocytes respond differently. Intriguingly, when iAstrocyte morphology, viability and marker expression was examined, we did not identify any modulatory effects of FUS^+MB^. To our knowledge this is the first study investigating the effects of FUS^+MB^ on patient-derived iAstrocytes of sporadic AD risk. We also examined the gene expression of several cytokines known to be secreted by astrocytes in basal conditions and following inflammatory stimulation [40-42]. Interestingly, our observations found some, although minimal effects of FUS^+MB^ on modulating iAstrocyte inflammatory responses. In particular, our results indicate that FUS^+MB^ may decrease the inflammatory response of *APOE3* astrocytes, with *IL-1β*, *IL-6* and *IL-8* expression reduced in *APOE3* iAstrocytes 24 h following FUS^+MB^ treatment with this response interestingly not observed in *APOE4* iAstrocytes. These results suggest that *APOE4* isoform carrying iAstrocytes might be less susceptible to potential immunomodulatory effects of FUS^+MB^. Our results demonstrated higher inflammatory cytokine expression in FUS^+MB^ treated *APOE4* iAstrocytes compared to their isogenic, *APOE3* converted pair, in-line with previous studies [74], potentially explaining why the reduction in inflammatory markers was not seen following FUS^+MB^. Although further experiments, such as assessment of cytokine secretion following FUS^+MB^, are needed to fully understand iAstrocyte response to FUS^+MB^, our results suggest that FUS^+MB^ does not elicit a drastic inflammatory response in iAstrocytes, with disease-specific differences remaining an important area of future investigation. Furthermore, with pericytes also playing a key inflammatory role in the BBB [75], it would be important to investigate their potential responses to FUS^+MB^. Indeed, the presence of all BBB cells, namely BECs, astrocytes and pericytes, might elicit responses not observed in monocultures, such as those used in this study.

Following characterization of FUS^+MB^ effects on iAstrocytes, we proceeded to establish iBEC+iAstrocyte co-cultures in an attempt to generate a more physiologically relevant BBB model for FUS^+MB^-mediated drug delivery screening. Consistent with previous studies [76, 77], co-culture of iBECs+iAstrocytes significantly increased barrier integrity in both *APOE3* and *APOE4* models. We then used the same parameters as for iBEC monocultures, to deliver therapeutic AD antibodies in the co-culture systems. Supporting the reproducibility of our delivery system, we were able to significantly increase the delivery of both Aducanumab and RNF5 using FUS^+MB^ in the *APOE3* and *APOE4* co-culture systems. Interestingly, however, we observed some differences between *APOE3* and *APOE4* models in the co-culture systems, not identified in iBEC monocultures. One was that Aducanumab delivery efficiency in the *APOE4* co-culture system was lower compared to *APOE3* co-cultures and *APOE4* iBEC monocultures. In contrast, RNF5 delivery efficiency was significantly higher in *APOE4* co-cultures compared to *APOE4* monocultures, although TEER in *APOE4* co-cultures was significantly higher. Without further investigation, it is difficult to hypothesize the physiological relevance of these observations in the human brain. One explanation could be that there are patient-specific differences in astrocyte function that modulate iBEC responses differently. Overall, the presence of astrocytes with iBECs is likely required to obtain a model that is more closely representative of the brain. Our established 2.5D BBB model further demonstrates the ability to culture iBECs and iAstrocytes in close contact within a supporting matrix and being able to use this model for FUS^+MB^-mediated drug delivery screening. Such a model, when fully optimized, will likely have higher translatability as a screening platform than traditional 2D models.

Overall, our data demonstrates a robust and reproducible BBB *in vitro* model to study FUS^+MB^-mediated therapeutic antibody delivery with the ability to identify potential disease- or patient-specific differences in treatment response. Whilst the iPSC-derived cells used in this model provide advantages in terms of modelling human disease phenotypes, one limitation is that they are prone to high levels of inter-cell line variation. Due to this, it might be difficult to accurately identify disease-specific differences, unless the number of biological replicates is increased. In addition, based on the *in vitro* results, it is difficult to predict how the increased antibody delivery following FUS^+MB^ would translate into the human brain and whether the observed increases in antibody fold change would be large enough to substantially increase therapeutic efficacy.

Finally, the physical dynamics of the free-floating MB in our system may differ from those observed in the constrained brain capillary, with the BBB opening *in vitro* being primarily achieved due to the formation of standing waves generated by the wave reflection from the media-air interphase (previously reported in [78]). In our system, with FUS transducer being positioned below the cell layer, the radiation force drives the MB from the pressure antinode of the standing wave towards the cell monolayer, facilitating interaction of oscillating MB and the cells [79, 80]. Although the FUS^+MB^ has been previously shown to achieve BBB opening in such a configuration *in vitro [18, 33]*, more complex model settings that incorporate all BBB cell types (BECs, astrocytes and pericytes) and potentially other CNS cell types as well as the ability to mimic blood flow and brain-blood interaction would be important for creating a more biologically and physically accurate model with higher translational capability. Finally, our model is currently limited to a fluorescence-based approach. Further improvements, such as by incorporating for example high-performance liquid chromatography (HPLC) could allow for a wider range of drugs to be assessed.

In conclusion, our reported model provides an important advancement in the field of ultrasound-based therapies, as there currently are no patient cell-based platforms to screen for FUS^+MB^-deliverable drugs or investigate the effects of FUS^+MB^ at the cellular level in humans. Using a patient-derived cell platform enables more rapid translation of findings into the clinic due to the ability to capture patient heterogeneity, a well-known occurrence amongst AD and other neurodegenerative disease patients. This platform can be used to screen for novel FUS-deliverable drugs that can then be tested in pre-clinical models and ultimately in patients. Finally, development of 2.5D and 3D BBB semi- or high-throughput models for FUS^+MB^ drug delivery screening will enable accelerated translation due to the ability to more closely mimic the human brain and brain cell interactions, downscale reagent use, and upscale the number of replicates.

## MATERIALS AND METHODS

### Experimental design

The objective of this research study was to develop an *in vitro* BBB-like model for FUS^+MB^-mediated therapeutic antibody delivery and investigate the contribution of the high-risk sporadic AD polymorphism *APOE4* allele on FUS^+MB^ response. With such cell platforms not existing, we hypothesized that by using a patient-derived cell model, we can accelerate the translation of FUS^+MB^ treatment for therapeutic drug delivery into the clinic by identifying FUS^+MB^-deliverable drugs. In addition, since the effects of FUS^+MB^ on a cellular and molecular level are not known, our aim was to carefully investigate these using microscopy, viability and RNA-seq analysis and identify potential differences between low risk (*APOE3*) and high risk (*APOE4*) AD cells. This was a controlled laboratory experiment, utilizing iBEC generation from n=3 *APOE3* carrier iPSC lines and n=3 *APOE4* carrier iPSC lines, which included one isogenic pair (*APOE4* converted to *APOE3*). In addition, n=2 *APOE3* and *APOE4* lines, including one isogenic pair were used to generate iAstrocytes for co-culture experiments. Cells in mono- and co-cultures were exposed to FUS^+MB^ to investigate the effects of the treatment on cell molecular responses and to investigate FUS^+MB^-mediated therapeutic antibody delivery. Sample size and experimental replicates were selected based on previous experiments with the current iPSC lines as well as based on what is commonly accepted in the field based on literature.

### Statistical analysis

Statistical analysis was performed using GraphPad Prism version 9.0.1. Data was analyzed using a two-tailed unpaired t-test with Welch’s correction with a *P* value of less than < 0.05 considered statistically significant. Values are shown as mean±SEM or mean±SD and specified in figure legends. The number of biological and independent replicates used for each experiment are specified in figure legends.

### Generation of human iPSC-derived brain endothelial-like cells and characterization of barrier integrity

Human iPSC lines were generated and characterized as previously described [27-29]. iPSCs were expanded on human recombinant vitronectin in StemFlex^TM^ medium (Thermo Fisher Scientific). iBEC differentiation was performed as previously described by us [18]. Forty-eight hours following purification of iBECs on collagen IV and fibronectin [18], barrier integrity of iBECs was characterized by measuring TEER using the EVOM3 Volt/Ohmmeter (World Precision Instruments) in 0.4 μm polyester or polycarbonate Transwell inserts (Sigma). Passive dextran permeability was measured by culturing iBECs on Ø 0.4 μm pore polyester or polycarbonate Transwell inserts and cells were exposed to 0.5 mg/mL fluorescein isothiocyanate (FITC)-conjugated 3 – 5 kDa dextran (Sigma) for 24h. The top and bottom well fluorescence was measured using a plate reader (Biotek Synergy H4) and clearance volume calculated as previously described [30].

### Human iPSC-derived iAstrocyte culture and establishment of co-cultures

For astrocyte differentiation, NPCs were first generated using STEMdiff^TM^ SMADi Neural Induction kit (Stemcell technologies) and generated neural progenitor cells were expanded using STEMdiff ^TM^ Neural Progenitor Medium. Following NPC characterization with Nestin and SOX2 (Fig. S5A), iAstrocyte differentiation was initiated by switching the medium to astrocyte medium (DMEM/F12+GlutaMAX, 1% N-2 supplement, 1% fetal bovine serum, all from Thermo Fisher Scientific) as previously described [81] and continued for at least 60 days. Astrocyte maturation was initiated 7 days prior to experiments by exposing cells to 10 ng/mL BMP-4 and CNTF as previously described [39]. Co-cultures were established by culturing iAstrocytes at density of 5,000 cells/cm^2^ in culture vessels in astrocyte medium supplemented with BMP4 and CNTF for 7 days. iBECs seeded in collagen IV and fibronectin coated Ø 3.0 μm Transwell inserts (Sigma) were placed in co-culture with astrocytes 24 h following purification and both cell types were maintained in human ESFM supplemented with B-27 (Thermo Fisher Scientific). Co-cultures were maintained for 24 h prior to experiments.

### Focused ultrasound (FUS) and microbubble (MB) treatments

In this study, a FUS system with a center frequency of 286 kHz was used. The system consisted of a transducer (Sonic Concepts) having an active diameter of 64 mm, with a 63.2 mm radius curvature, housed in a 82 mm spherical shell with a central opening of Ø 20 mm used with the RF amplifier (Electronics & Innovation, Ltd). The focus of the transducer had dimensions of 6.04 mm × 39.49 mm (Ø focal width x focal length). Transducer was mounted in a custom-made plexiglass holder and immersed in a water bath filled with de-gassed water. Transducer pressure wave calibration was performed using needle hydrophone. All experiments presented in this study were performed using the following FUS settings: 286 kHz center frequency, 0.3 MPa peak rarefactional pressure applied outside of the cell culture plate, 50 cycles/burst, burst period 20 ms, and a 120-s sonication time. Prior to FUS treatment, cells were exposed to phospholipid-shelled microbubbles with octafluoropropane gas core, prepared in-house, following previously described chemical synthesis protocol [82]. For dextran and therapeutic delivery studies, iBECs were seeded on Ø 3.0 μm pore Transwells at 1.4×10^6^ cells/cm^2^. The effect of FUS^+MB^ was tested on iBECs 48– 72h after subculture on Transwell inserts. FITC-conjugated dextran (150 kDa) was added at 0.5 mg/ml and AlexaFluor™-647-conjugated anti-amyloid-β (Aducanumab) and anti-Tau (RNF5, [26]) therapeutic antibodies were added at 1 μM. MB (10 μL per Transwell) were then added to the wells aseptically directly before the FUS treatment. Cells were then exposed to FUS and 24 h after FUS treatment, media samples from top and bottom chamber of the Transwell were collected for spectrofluorometric analysis. Fluorescence of dextran was measured at 490 nm excitation/520 nm emission and fluorescence of antibodies was measured at 633 nm excitation/665 nm emission using a plate reader (Biotek Synergy H4). Clearance volume in UT and FUS^+MB^ treated Transwells was calculated as previously described [30] and data presented as fold change to UT.

### Lactate dehydrogenase (LDH) cytotoxicity assay

To assess the effects of FUS^+MB^ on iBEC and iAstrocyte viability, cell culture media samples were collected 1 h and 24 h after FUS^+MB^ exposure and stored in -80 °C until analysis. The level of lactate dehydrogenase (LDH) enzyme in the collected media was determined using CyQUANT LDH Cytotoxicity Assay (Thermo Fisher Scientific) following manufacturer’s instructions. Absorbance was measured at 490 nm and 680 nm using a plate reader. To determine LDH activity, the 680 nm absorbance value (background) was subtracted from the 490 nm absorbance and compared between treatment conditions.

### RNA extraction, cDNA synthesis and quantitative real-time PCR

For RNA collection, cells were rinsed with PBS, exposed to TRIzol^TM^ reagent and scraped off the culture plate using a pipette tip. Total RNA was extracted using the Direct-zol RNA Miniprep Kit (Zymo Research) according to the manufacturer’s instructions and treated in-column with DNase I. RNA quality and quantity was measured using NanoDrop^TM^ Spectrophotometer, after which RNA was converted to cDNA using SensiFAST^TM^ cDNA synthesis kit (Bioline) according to the manufacturer’s instructions. The qPCR run was performed as triplicate for each sample on QuantStudio^TM^ 5 Real-Time PCR system with run conditions as follows: 2 min at 95°C followed by 40 cycles of 5 s at 95°C and 30 s at 60°C. Ct values were normalised to Ct values of 18S endogenous control (ΔCt values), which was found to be consistent across cell lines, conditions and timepoints. ΔΔCt values were calculated as 2^(-ΔCt)^ and presented as ΔΔCt multiplied by 10^6^ or as fold change. Primer sequences used in this study are presented in Table S3.

### Bulk RNA-sequencing (RNA-seq)

For transcriptome analysis of FUS^+MB^ treated iBECs, bulk RNA-seq was performed. Briefly, iBECs were purified on collagen IV and fibronectin coated 24-well plates, then treated with 20 μL of MB followed by FUS with parameters described above. RNA samples were harvested at 1 h and 24 h timepoints from untreated and FUS^+MB^ treated cells. Total RNA was extracted as described above. The quality of total RNA was determined using the Agilent TapeStation system. In total, 24 samples underwent RNA-seq: samples from 6 patients (n = 3 *APOE3* and n = 3 *APOE4*) each with FUS^+MB^ treatment or UT at both 1 h and 24 h timepoints.

Library preparation was performed using Illumina TruSeq Stranded mRNA kit and libraries were sequenced using the Illumina NextSeq 550 platform at the QIMR Berghofer Next Generation Sequencing facility with 75 bp reads sequenced to ∼ 40 million reads per sample. Sequence reads were trimmed for adaptor sequences using Cutadapt version 1.9 [83] and aligned using STAR version 2.5.2a [84] to the human GRCh37 assembly with the gene, transcript, and exon features of Ensembl (release 70) gene model. Quality control metrics were computed using RNA-SeQC version 1.1.8 [85] and expression was estimated using RSEM version 1.2.30 [86].

All downstream RNA-seq analysis was performed using R version 3.6.2. Differential expression analysis was performed using edgeR’s quasi-likelihood pipeline version 3.28.0 [87-89]. Specifically, only protein-coding genes that passed the minimum expression filter using edgeR’s filterByExpr function with default settings were kept for further analysis. Two design matrices were constructed for the analyses herein. For the comparisons between FUS^+MB^ treatment vs UT at different time points irrespective of genotype, an additive linear model was used, which incorporated a patient term and the remaining experimental conditions combined into one factor. Specifically, we used model.matrix(∼Patient + Treatment.Time), where Patient was one of 6 patient IDs, Treatment is UT or FUS^+MB^, and Time is 1 h or 24 h. For the comparisons between genotypes, all experimental conditions were combined into a single factor. Specifically, we used model.matrix(∼Genotype.Treatment.Time), where Genotype is *APOE3* or *APOE4*, and Treatment and Time as per above. The glmQLFit() function was used to fit a quasi-likelihood negative binomial generalized log-linear model to the read counts for each gene. Using the glmQLFTest() function, we conducted gene-wise empirical Bayes quasi-likelihood F-tests for a given contrast. Differentially expressed genes (DEGs) were determined using a false discovery rate (FDR) < 0.05. To perform gene ontology (GO) term analysis, multiple functions from the clusterProfiler package version 3.14.3 were utilized. First, the bitr function was used to convert gene IDs of DEGs from Ensembl to Entrez. Entrez IDs were subsequently passed to the enrichGO function, before plotting the results with the dotplot function [90].

### Immunofluorescence

For immunofluorescence characterization, iBECs grown on collagen IV and fibronectin, washed with PBS and fixed with ice cold 100% methanol for 5 min or 4% paraformaldehyde (PFA) for 15 min. Astrocytes were fixed with 4% PFA. Following PFA fixing, cells were permeabilized for 10 min with 0.3 % Triton-X. Cells were then blocked for 1 – 2 h at RT with 2% bovine serum albumin and 2% normal goat serum in PBS. Primary antibodies (Table S4) were diluted in blocking solution and incubated overnight at 4°C. The following day, cells were washed three times with PBS, then incubated with secondary antibodies (Table S4) diluted in blocking solution for 1 – 2 h at RT in the dark. Finally, cells were washed three times with PBS, Hoechst counterstain was performed, and coverslips were mounted with ProLong Gold Antifade (Thermo Fisher Scientific). Images were obtained at 10X or 20X magnification using a Zeiss 780 confocal microscope. Image brightness was increased for presentation purposes using ImageJ.

### 2.5D BBB-like model

The 2.5D BBB-like model was established using photo-crosslinkable synthetic LunaGel^TM^ (Gelomics, Brisbane, Australia). For this, iAstrocytes were first seeded in the gel prior to iBEC seeding. Briefly, iAstrocytes were differentiated for approximately 60 days, after which they were detached from their culture vessel with TrypLE (Thermo Fisher Scientific) and a cell count was performed. iAstrocytes were then centrifuged to a pellet (300 x g, 5 min) and resuspended in LunaGel^TM^ mixed 1:1 with photoinitiator as per manufacturer’s instructions. iAstrocytes were then seeded in a clear 96-well plate at a ratio of 10,000 cells per 50 μL of gel per well. The gel was polymerized for 30 s using the LunaCrosslinker^TM^ to ensure a soft gel for iAstrocyte proliferation. Following polymerization, 100 μL of astrocyte medium was added on top of the gel. The following day cells were supplemented with BMP-4 and CNTF for 7 days after which seeding of iBECs was performed. Prior to seeding iBECs on the iAstrocyte containing LunaGel^TM^, iBECs were purified on a collagen IV and fibronectin coated culture flask for 24 h. Before iBEC seeding, astrocyte medium was removed, and a thin layer (25 μL) of high stiffness LunaGel^TM^ was seeded on top of the iAstrocyte layer and polymerized for 60 s. The layer of high stiff gel was then coated with collagen IV and fibronectin for 1 h prior to seeding of iBECs. iBECs were detached using TrypLE and a cell count was performed. iBECs were resuspended in ESFM + B27 supplemented with 10 μM retinoic acid, 10 μM ROCKinhibitor and 1.3 μM as previously described [46, 47]. The coating solution was removed from the gel and iBECs seeded at 150,000 cells per well on top of the iAstrocyte gel. iBECs were allowed to attach for 24h, after which the medium was switched to ESFM+B27 with no supplementation. Aducanumab delivery using FUS^+MB^ was performed as described above. Following 24 h, the supernatant was removed and replaced with PBS. Antibody fluorescence intensity was measured in the gel using a Biotek synergy H4 plate reader as described above.

## Acknowledgments

We thank the QIMR Berghofer Next Generation Sequencing facility for sample processing and RNA-seq as well as providing technical advice. We thank R. Koufariotis for RNA-seq data alignment and N. Waddell for assisting with analysis and workflow design of RNA-seq results. We thank J. Pearson and the Genome Informatics Group for data management and computational support.

## Abbreviations

AJ: adherens junction
APOE: apolipoprotein E
AQP4: aquaporin-4
Aβ: amyloid-β
BBB: blood-brain barrier
BEC: brain endothelial cell
BMP-4: bone morphogenic protein 4
CCL2: C-C motif chemokine ligand 2
CNTF: ciliary neurotrophic factor
DEG: differentially expressed gene
ESFM: endothelial serum-free medium
FDR: false discovery rate
FUS: focused ultrasound
GFAP: glial fibrillary acidic protein
hiPSC: human induced pluripotent stem cell
HPLC: high-performance liquid chromatography
iBEC: induced brain endothelial-like cell
IL: interleukin
LDH: lactate dehydrogenase
MB: microbubble
NPC: neural progenitor cell
p-Tau: phosphorylated Tau
PCA: principal component analysis
RNA-seq: RNA sequencing
TEER: trans-endothelial electrical resistance
TJ: tight junction
UT: untreated

## Funding

This work was supported by:

National Health and Medical Research Council (NHMRC) Senior Research Fellowship (1154389) (AP)

National Health and Medical Research Council (NHMRC) Senior Research Fellowship (1118452) (ARW)

DH was supported by Yulgilbar Alzheimer’s Research Program grant

JMW is a recipient of The University of Queensland PhD scholarship

## Author contributions

Conceptualization: JMW, ARW, LEO

Methodology: JMW, JCSC, RLJ, LAM, DH, LC, GL, JS, WL, RMN, LEO

Investigation: JMW, JCSC, RLJ, LEO

Visualization: JMW, RLJ, LEO

Funding acquisition: AP, JG, ARW

Project administration: JG, ARW

Supervision: RMN, AP, JG, ARW, LEO

Writing – original draft: JMW, RLJ, ARW, LEO

Writing – review & editing: JG

## Competing interests

The authors have declared that no competing interest exists.

## Data and materials availability

RNA sequencing metadata that support the findings of this study are openly available in the European Genome-Phenome Archive (EGA) at EGA Accession Number: EGAS00001005944. All other data are presented in the main text or the supplementary materials or can be made available by contacting the corresponding author.

---

**Fig. S1.**
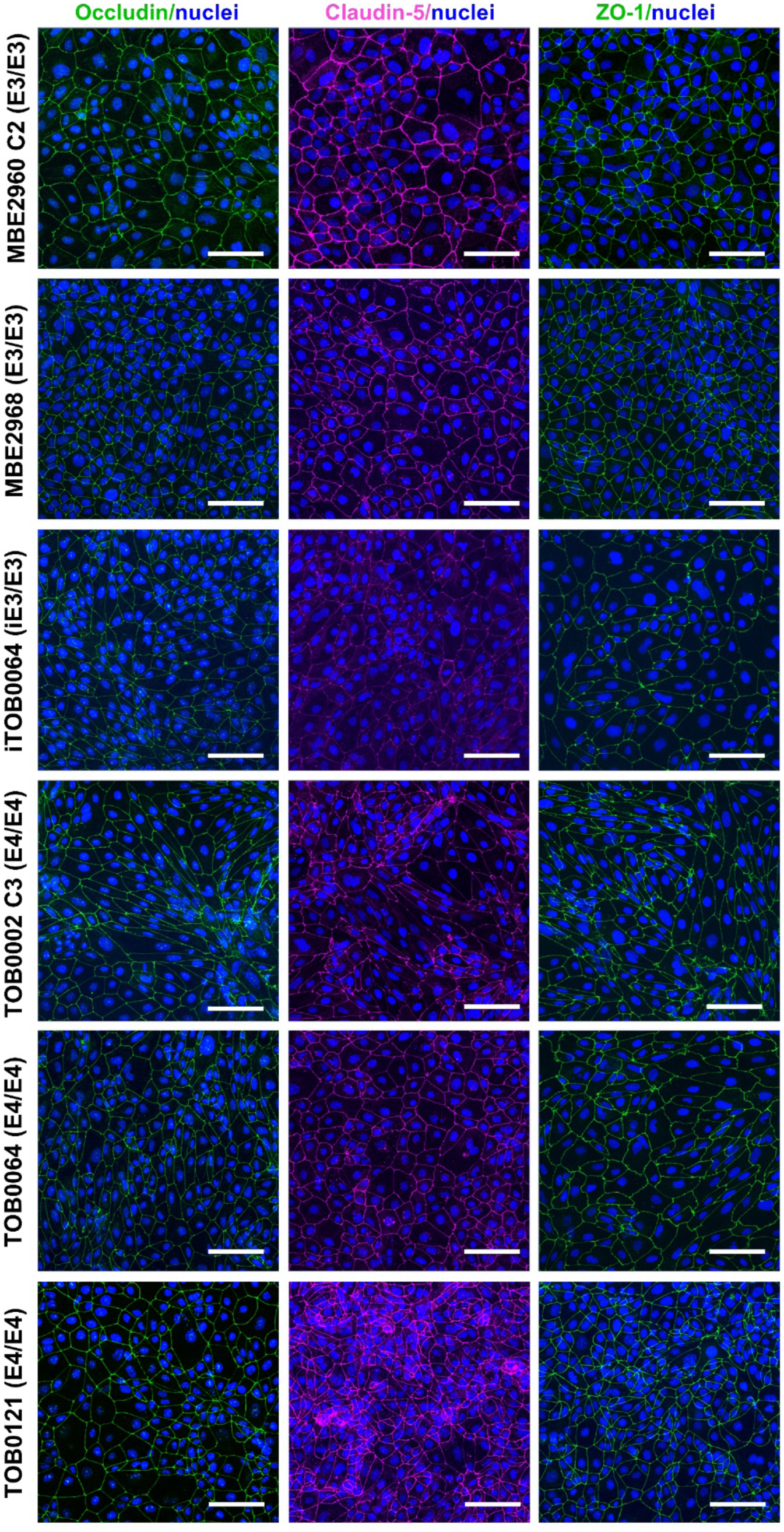
Characterization of iBECs generated from individual iPSC lines. Immunofluorescence images of occludin (green), claudin-5 (magenta) and ZO-1 (green) in iBECs generated from *APOE3* and *APOE4* induced pluripotent stem cells (iPSCs) used in this study. Hoechst counterstain, scale bar = 100 μm.

**Fig. S2.**
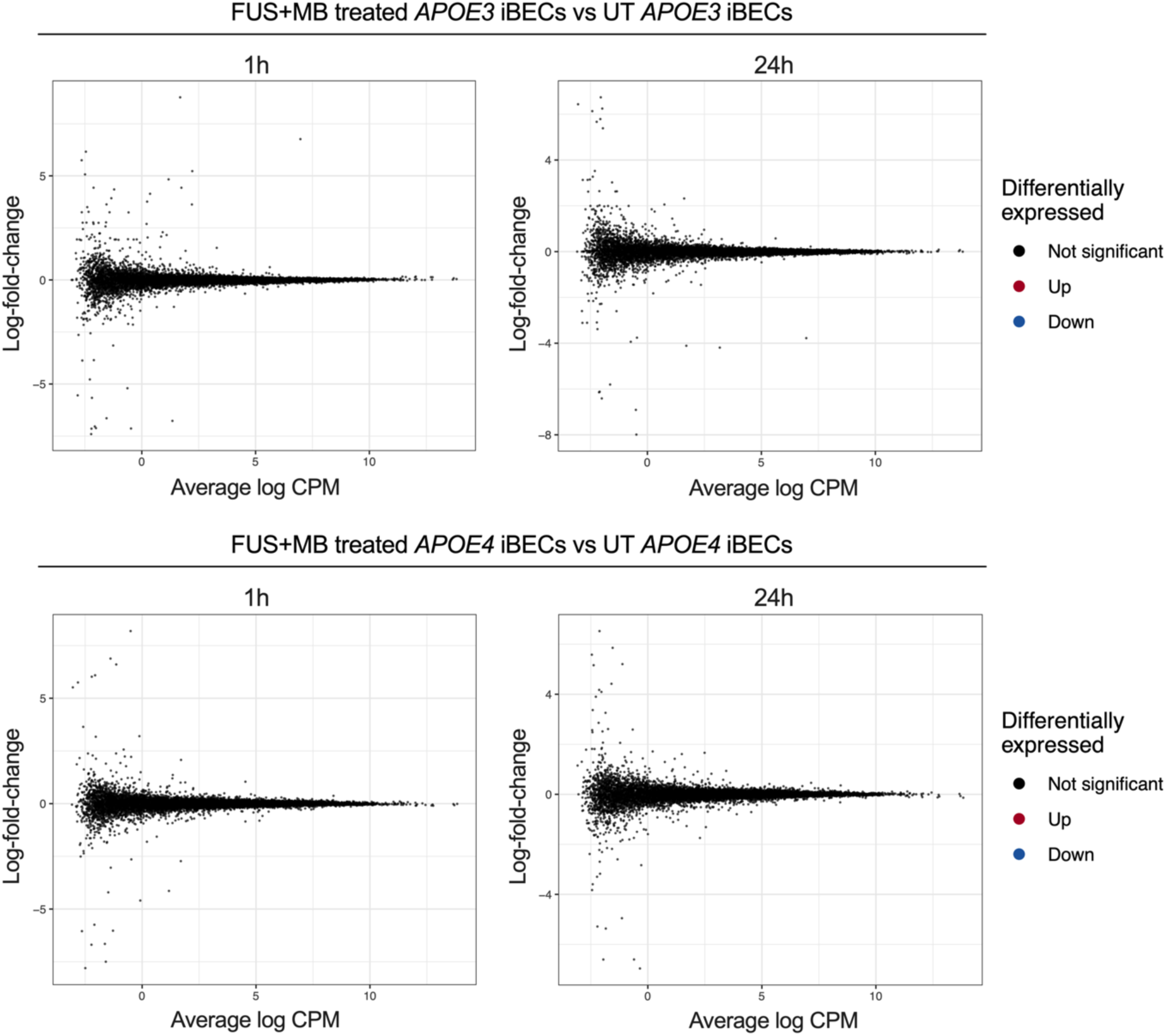
Differential expression analysis of FUS+MB treated iBEC vs untreated iBECs at 1 h and 24 h for APOE3 and APOE4 genotypes. Mean-difference (MD) plots showing log-fold-change versus average log expression values (log2 counts per million, CPM). Upper panel: FUS+MB treated APOE3 iBECs at 1 h vs untreated (UT) APOE3 iBECs at 1 h and FUS+MB treated APOE3 iBECs at 24 h vs UT APOE3 iBECs at 24 h. Lower panel: FUS+MB treated APOE4 iBECs at 1 h vs UT APOE4 iBECs at 1 h and FUS+MB treated APOE4 iBECs at 24 h vs UT APOE4 iBECs at 24 h.

**Fig. S3.**
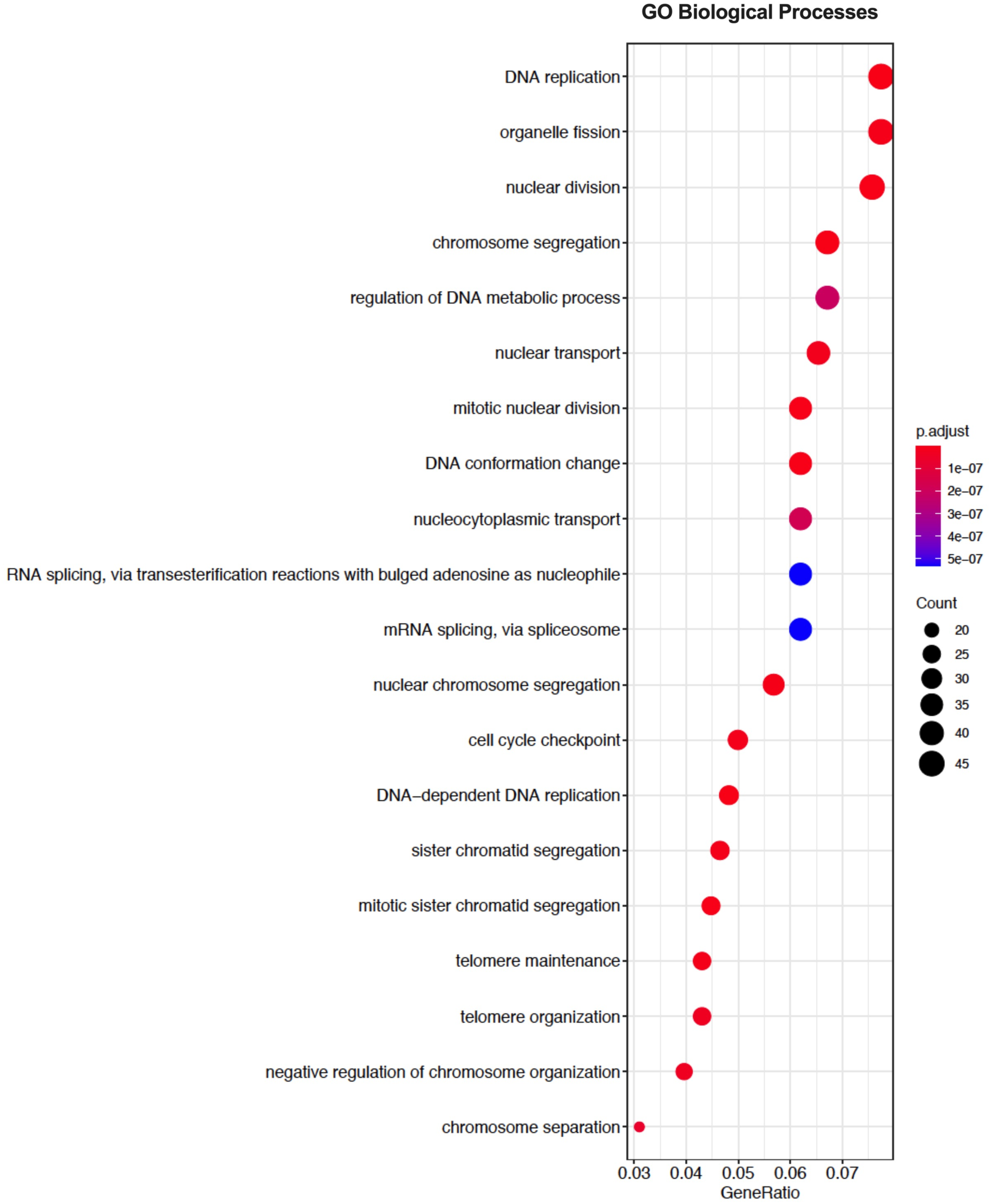
Dotplot of top gene ontology (GO) terms from sub ontology Biological Process enriched from comparison of UT *APOE4* iBEC at 1 h vs *APOE3* iBEC at 1 h. The top 20 GO processes according to p-value plotted in order of gene ratio. The size of the dots represents the number of genes associated with the GO term and the color of the dots represent the *P*-adjusted values. The differentially expressed genes (FDR <0.05) were used for analysis.

**Fig. S4.**
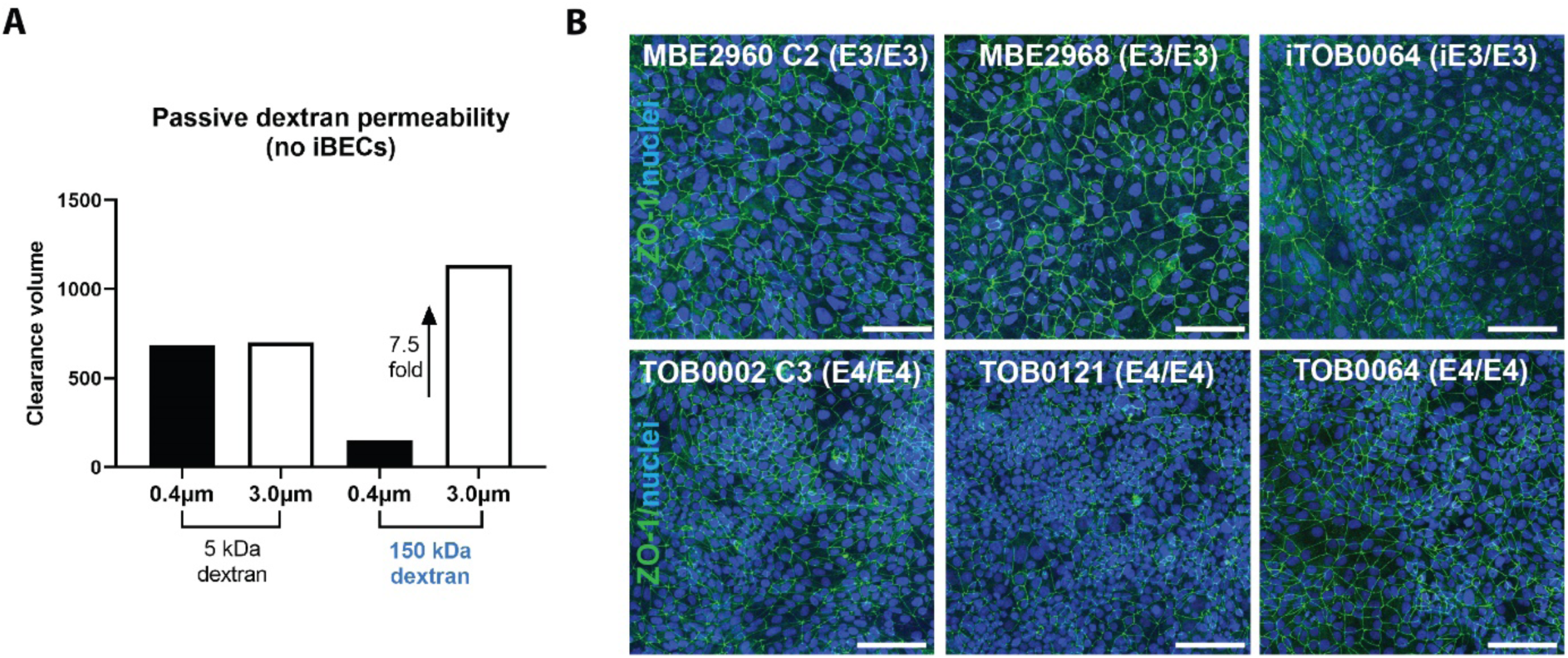
Optimization of the Ø 3.0 μm pore Transwell model. (**A**) Passive permeability (clearance volume) of 5 kDa and 150 kDa dextran in collagen IV and fibronectin coated (no iBEC containing) 0.4 μm and 3.0 μm pore Transwell inserts. (**B**) Immunofluorescence of ZO-1 (green) in each individual *APOE3* and *APOE4* iBEC line seeded on 3.0 μm pore Transwell inserts (Hoechst counterstain, scale bar = 100 μm).

**Fig. S5.**
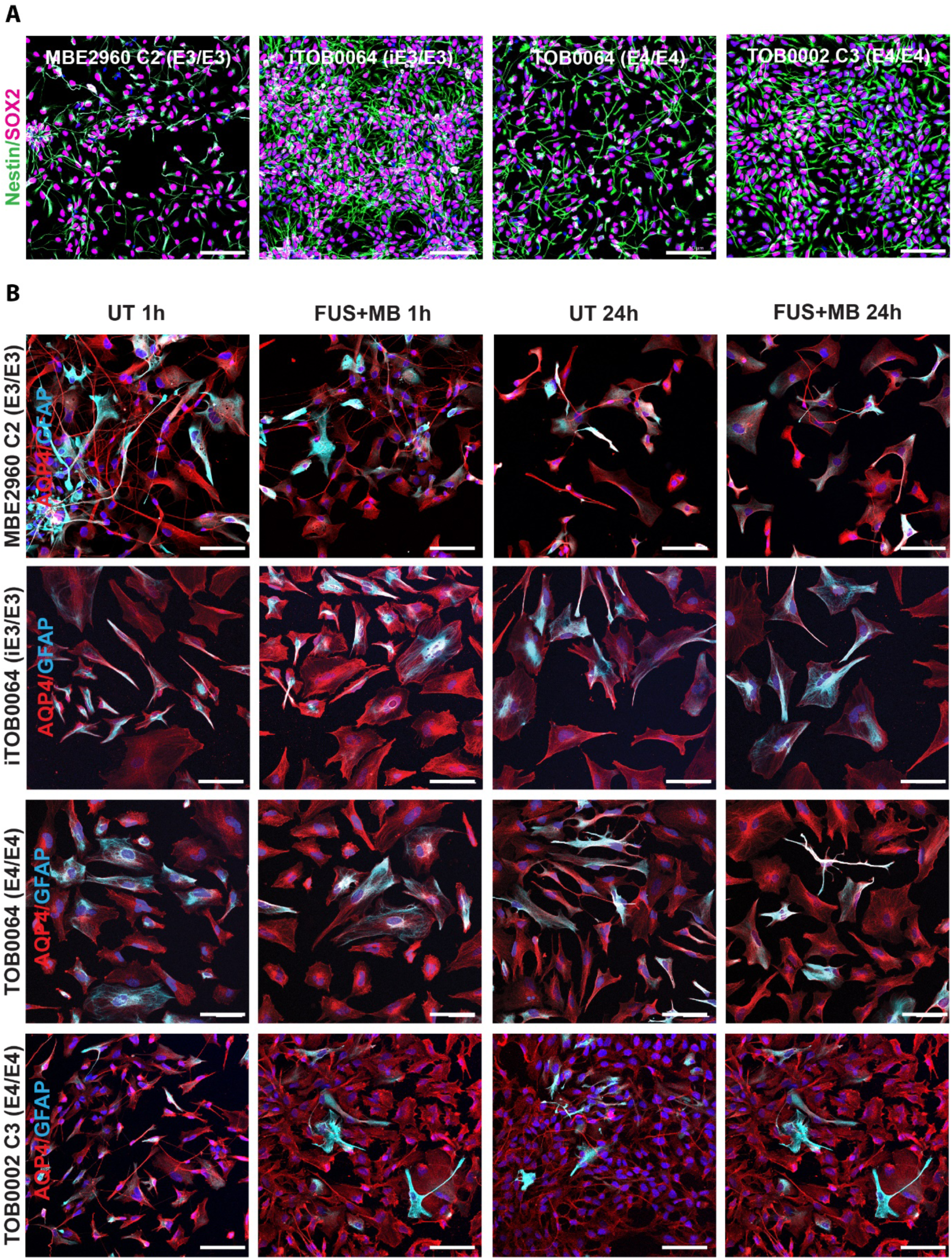
Neural progenitor cell generation and FUS^+MB^ treatment of iAstrocytes. (**A**) Immunofluorescence of nestin (green) and SOX2 (magenta) in neural progenitor cells (NPCs) generated from *APOE3* and *APOE4* induced pluripotent stem cells (iPSCs) used in this study. (**B**) Immunofluorescence of *APOE3* and *APOE4* iAstrocytes generated from individual iPSC lines in untreated (UT) and focused ultrasound + microbubble (FUS^+MB^) conditions 1 h and 24 h following treatment stained with AQP4 (red) and GFAP (cyan) (Hoechst counterstain, scale bar = 100 μm).

**Fig. S6.**
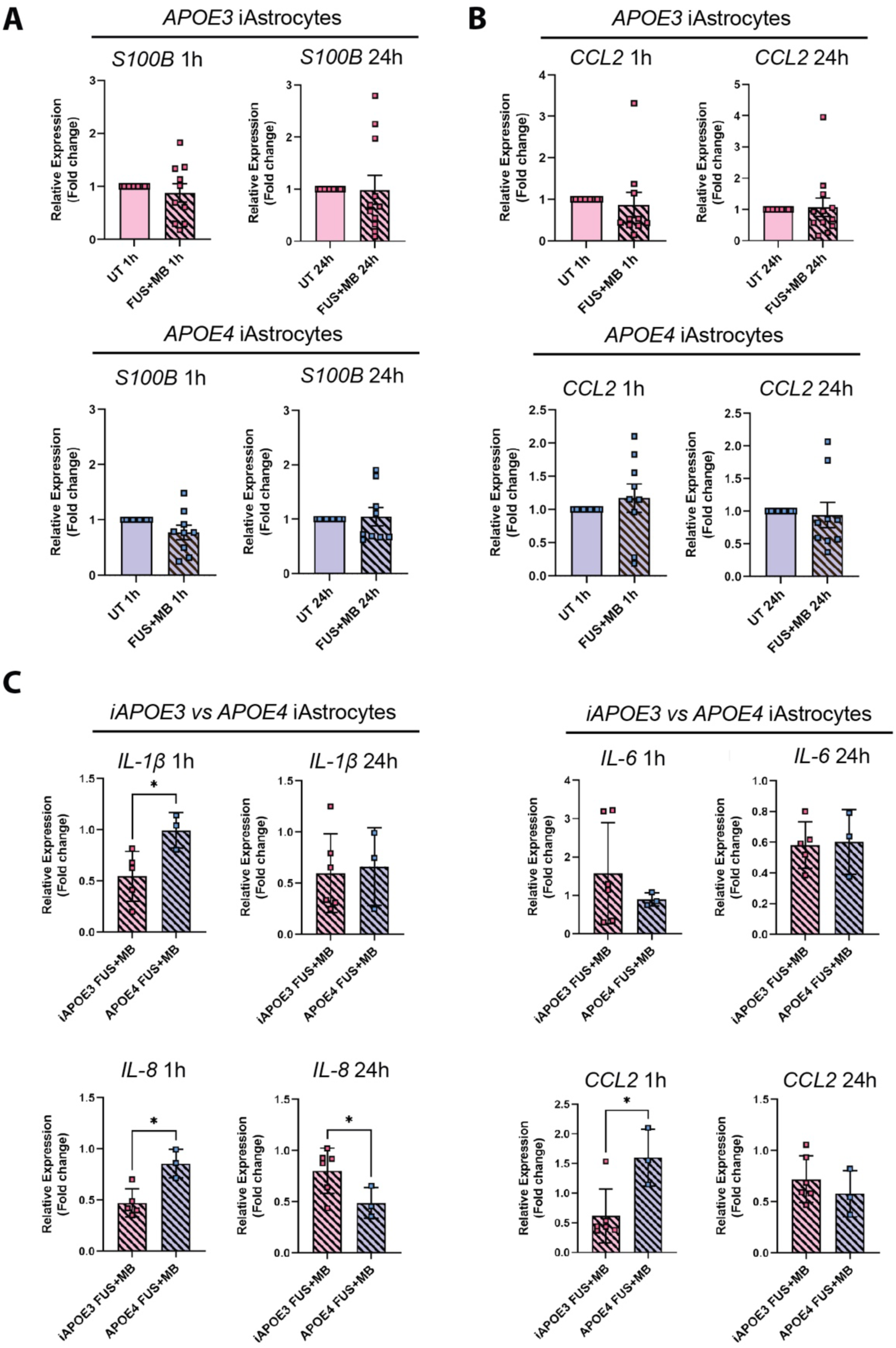
Effect of FUS^+MB^ on iAstrocyte gene expression. Relative gene expression (fold change) of (**A**) astrocyte marker *S100B* and (**B**) inflammatory cytokine *CCL2* in UT and FUS^+MB^ treated *APOE3* and *APOE4* iAstrocytes 1 h and 24 h after treatment, error bars = SEM. (**C**) Comparison of relative gene expression (fold change) of inflammatory markers between an isogenic *iAPOE3* and *APOE4* iAstrocyte pair at 1 h and 24 h after FUS^+MB^ treatment, error bars = SD. \**P*<0.05 by Unpaired t-test with Welch’s correction.

## SUPPLEMENTARY MATERIALS

**Table S1. Results from differential expression analysis of UT APOE4 iBECs compared to UT APOE3 iBECs at 1 h.** Genes sorted by P-value. Differentially expressed genes (DEGs) defined by FDR < 0.05.

Table can be found in the .xlsx format

**Table S2. Results from gene ontology (GO) enrichment analysis of sub ontology Biological Process from comparison of UT *APOE4* iBEC at 1 h vs *APOE3* iBEC at 1 h**.

Table can be found in the .xlsx format

**Table S3.**
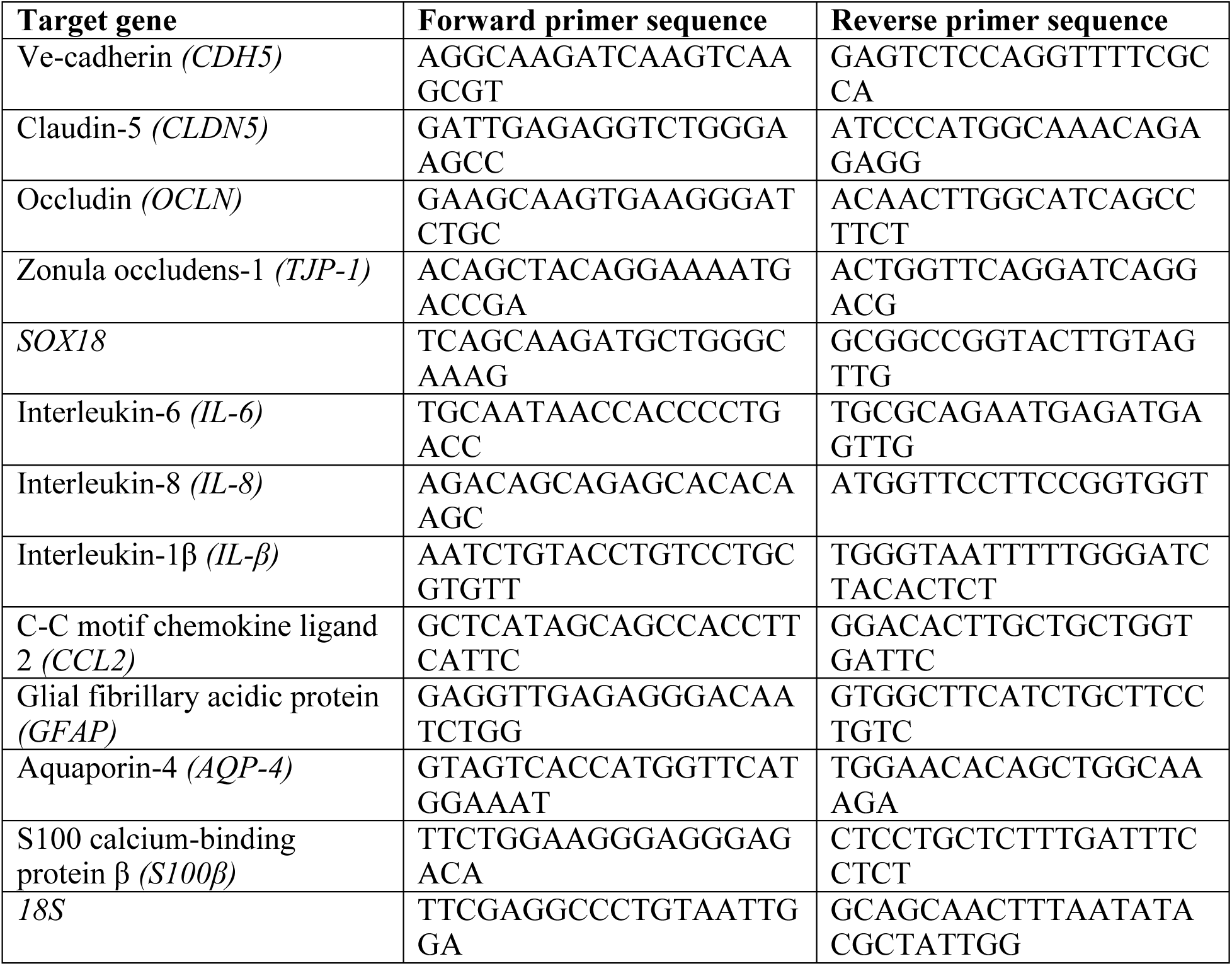
Primer sequences used in the study.

**Table S4.** Antibodies used in the study.

| **Primary antibodies** | **Species** | **Source** | **Identifier** |
| --- | --- | --- | --- |
| ZO-1 | mouse | Invitrogen | Cat# 339100 |
| occludin | rabbit | Invitrogen | Cat# 711500 |
| claudin-5 | mouse | Invitrogen | Cat# 352500 |
| aquaporin-4 | mouse | Abcam | Cat# ab9512 |
| GFAP | rabbit | Agilent/ Dako | Cat# Z0334 |
| nestin | mouse | Abcam | Cat# ab22035 |
| SOX2 | rat | Invitrogen | Cat# 14981182 |
| **Secondary antibodies** | **Species** | **Source** | **Identifier** |
| anti-mouse Alexa Fluor 488 | goat | Invitrogen | Cat# A11029 |
| anti-mouse Alexa Fluor 594 | goat | Invitrogen | Cat# A11032 |
| anti-mouse Alexa Fluor 647 | goat | Invitrogen | Cat# A32728 |
| anti-rabbit Alexa Fluor 488 | goat | Invitrogen | Cat# A11034 |
| anti-rat Alexa Fluor 647 | goat | Invitrogen | Cat# A21247 |

## Notes

### Competing Interest Statement

The authors have declared no competing interest.

